# Structural basis of a public antibody response to SARS-CoV-2

**DOI:** 10.1101/2020.06.08.141267

**Authors:** Meng Yuan, Hejun Liu, Nicholas C. Wu, Chang-Chun D. Lee, Xueyong Zhu, Fangzhu Zhao, Deli Huang, Wenli Yu, Yuanzi Hua, Henry Tien, Thomas F. Rogers, Elise Landais, Devin Sok, Joseph G. Jardine, Dennis R. Burton, Ian A. Wilson

**Affiliations:** Department of Integrative Structural and Computational Biology, The Scripps Research Institute, La Jolla, CA 92037, USA; Department of Immunology and Microbiology, The Scripps Research Institute, La Jolla, CA 92037, USA; IAVI Neutralizing Antibody Center, The Scripps Research Institute, La Jolla, CA 92037, USA; Consortium for HIV/AIDS Vaccine Development (CHAVD), The Scripps Research Institute, La Jolla, CA 92037, USA; Division of Infectious Diseases, Department of Medicine, University of California, San Diego, La Jolla, CA 92037, USA; IAVI, New York, NY10004, USA; Ragon Institute of Massachusetts General Hospital, Massachusetts Institute of Technology, and Harvard University, Cambridge, MA 02139, USA; The Skaggs Institute for Chemical Biology, The Scripps Research Institute, La Jolla, CA, 92037, USA

**Author notes:** These authors contributed equally to this work (Meng Yuan, Hejun Liu, Nicholas C. Wu). Correspondence (I.A.W.).

## Abstract

Molecular-level understanding of human neutralizing antibody responses to SARS-CoV-2 could accelerate vaccine design and facilitate drug discovery. We analyzed 294 SARS-CoV-2 antibodies and found that IGHV3-53 is the most frequently used IGHV gene for targeting the receptor binding domain (RBD) of the spike (S) protein. We determined crystal structures of two IGHV3-53 neutralizing antibodies +/- Fab CR3022 ranging from 2.33 to 3.11 Å resolution. The germline-encoded residues of IGHV3-53 dominate binding to the ACE2 binding site epitope with no overlap with the CR3022 epitope. Moreover, IGHV3-53 is used in combination with a very short CDR H3 and different light chains. Overall, IGHV3-53 represents a versatile public VH in neutralizing SARS-CoV-2 antibodies, where their specific germline features and minimal affinity maturation provide important insights for vaccine design and assessing outcomes.

## MAIN

The ongoing COVID-19 pandemic, which is caused by severe acute respiratory syndrome coronavirus 2 (SARS-CoV-2), is far from an end (*1*). The increasing global health and socioeconomic damage require urgent development of an effective COVID-19 vaccine. While multiple vaccine candidates have entered clinical trials (*2*), the molecular features that contribute to an effective antibody response are not clear. Over the past decade, the concept of a public antibody response (also known as multidonor class antibodies) to specified microbial pathogens has emerged. A public antibody response describes antibodies that have shared genetic elements and modes of recognition, and can be observed in multiple individuals against a given antigen. Such responses to microbial pathogens have been observed against influenza (*3*), dengue (*4*), malaria (*5*), and HIV (*6*). Identification of public antibody responses and characterization of the molecular interactions with cognate antigen can provide insight into the fundamental understanding of the immune repertoire and its ability to quickly respond to novel microbial pathogens, as well as facilitate rational vaccine design against these pathogens (*7, 8*).

The spike (S) protein is the major surface antigen of SARS-CoV-2. The S protein utilizes its receptor-binding domain (RBD) to engage the host receptor ACE2 for viral entry (*9–12*). Therefore, RBD-targeting antibodies could neutralize SARS-CoV-2 by blocking ACE2 binding. A number of antibodies that target the RBD of SARS-CoV-2 have now been discovered in very recent studies (*13–28*). We compiled a list of 294 SARS-CoV-2 RBD-targeting antibodies where information on IGHV gene usage is available (*17–28*) (Table S2), and found that IGHV3-53 is the most frequently used IGHV gene among such antibodies (Fig. 1A). Of 294 RBD-targeting antibodies, 10% are encoded by IGHV3-53, as compared to only 0.5% to 2.6% in the repertoire of naïve healthy individuals (*29*) with a mean of 1.8% (*30*). The prevalence of IGHV3-53 in the antibody response in SARS-CoV-2 patients has also been recognized in some recent antibody studies (*20, 22, 27*). These observations indicate that IGHV3-53 represents a frequent and public antibody response to the SARS-CoV-2 RBD.

**Figure 1.**
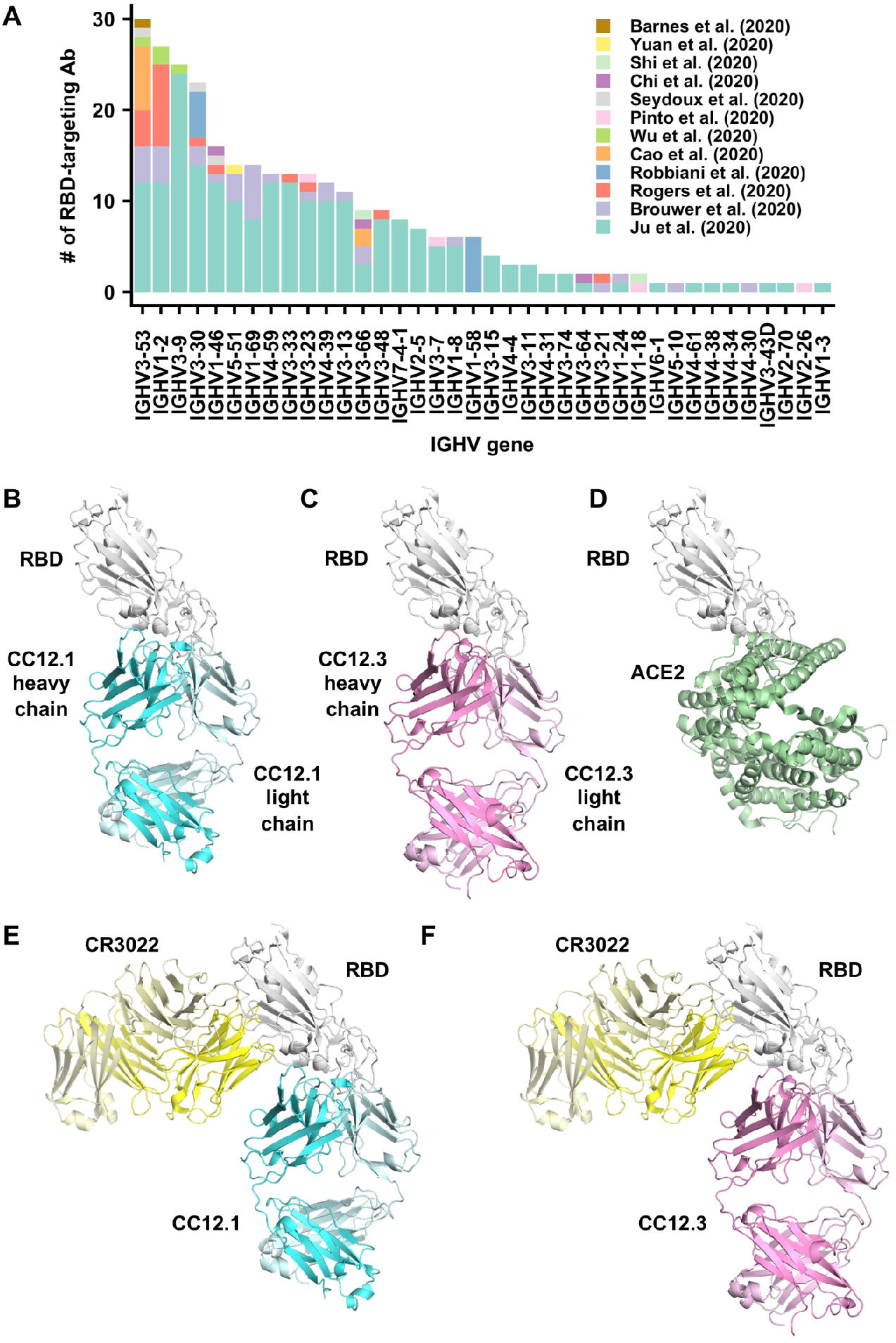
Structures of two IGHV3-53 antibodies. **(A)** The distribution of IGHV gene usage is shown for a total of 294 RBD-targeting antibodies (*17–28*). **(B-F)** Crystal structures of **(B)** CC12.1 in complex with SARS-CoV-2 RBD, **(C)** CC12.3 with SARS-CoV-2 RBD, **(D)** human ACE2 with SARS-CoV-2 RBD (PDB 6M0J) (*12*), **(E)** SARS-CoV-2 RBD with CC12.1 and CR3022, and **(F)** SARS-CoV-2 RBD with CC12.3 and CR3022.

To understand the molecular features that endow IGHV3-53 with the ability to act as a public antibody, we determined crystal structures of two IGHV3-53 neutralizing antibodies, namely CC12.1 and CC12.3, in complex with the SARS CoV-2 RBD and also in the presence of the SARS-CoV1/2 cross-reactive Fab CR3022 (*17*). CC12.1 and CC12.3 were previously isolated from a SARS-CoV-2-infected patient and shown to be SARS-CoV-2 RBD-specific (*27*). Although CC12.1 and CC12.3 are both encoded by IGHV3-53, CC12.1 utilizes IGHJ6, IGKV1-9, and IGKJ3, whereas CC12.3 utilizes IGHJ4, IGKV3-20, and IGKJ1. This variation in IGHJ, IGKV, and IGKJ usage indicates that CC12.1 and CC12.3 belong to different clonotypes, but are encoded by a common IGHV3-53 germline gene. IgBlast analysis (*31*) shows that IGHV and IGKV of CC12.1 are only 1% somatically mutated at the nucleotide sequence level (two amino-acid changes each). Similarly, the IGHV and IGKV of CC12.3 are also minimally somatically mutated at 1.4% in both IGHV (four amino-acid changes) and IGKV (a single amino-acid deletion). The binding affinities (K_d_) of Fabs CC12.1 and CC12.3 to SARS-CoV-2 RBD are 17 nM and 14 nM, respectively (Fig. S2). Moreover, competition experiments suggest that CC12.1 and CC12.3 bind to a similar epitope, which overlaps with the ACE2 binding site, but not the CR3022 epitope (Fig. S3).

We determined four complex crystal structures, CC12.1/RBD, CC12.3/RBD, CC12.1/RBD/CR3022, and CC12.3/RBD/CR3022 at resolutions of 3.11 Å, 2.33 Å, 2.90 Å, and 2.70 Å, respectively (Table S1). CC12.1 and CC12.3 bind to the ACE2 binding site on SARS-CoV-2 RBD with an identical angle of approach (Fig. 1B-F). Interestingly, another IGHV3-53 antibody B38, whose structure was determined recently (*23*), also binds to the ACE2 binding site on SARS-CoV-2 RBD in a similar manner (Fig. S4). Similar to the ACE2 binding site (*11*), the epitopes of these antibodies can only be accessed when the RBD is in the “up” conformation (Fig. S5). Among 16 ACE2 binding residues on RBD, 10 are within the epitopes of CC12.1 and B38, and 6 are in the epitope of CC12.3 (Fig. 2A-D). Many of the epitope residues are not conserved between SARS-CoV-2 and SARS-CoV (Fig. 2E), explaining their lack of cross-reactivity (*27*). The buried surface area (BSA) from the heavy-chain interaction is quite similar in CC12.1 (723 Å^2^), CC12.3 (698 Å^2^), and B38 (713 Å^2^). In contrast, the light-chain interaction is much smaller for CC12.3 (176 Å^2^) compared to CC12.1 (566 Å^2^) and B38 (495 Å^2^), consistent with different light-chain gene usage. While both CC12.1 and B38 utilize IGKV1-9, CC12.3 utilizes IGKV3-20. This observation suggests that IGHV3-53 can pair with different light chains to target the ACE2 binding site of the SARS-CoV-2 RBD. Given that CC12.3 (80% BSA from the heavy chain) binds the RBD with similar affinity to CC12.1 (56% BSA from heavy chain) (Fig. S2), the light-chain identity seems not to be as critical as the heavy chain. In fact, among the RBD-targeting IGHV3-53 antibodies, nine different light chains are observed, although IGKV1-9 and IGKV3-20 are the most frequently found to date (Fig. S6).

**Figure 2.**
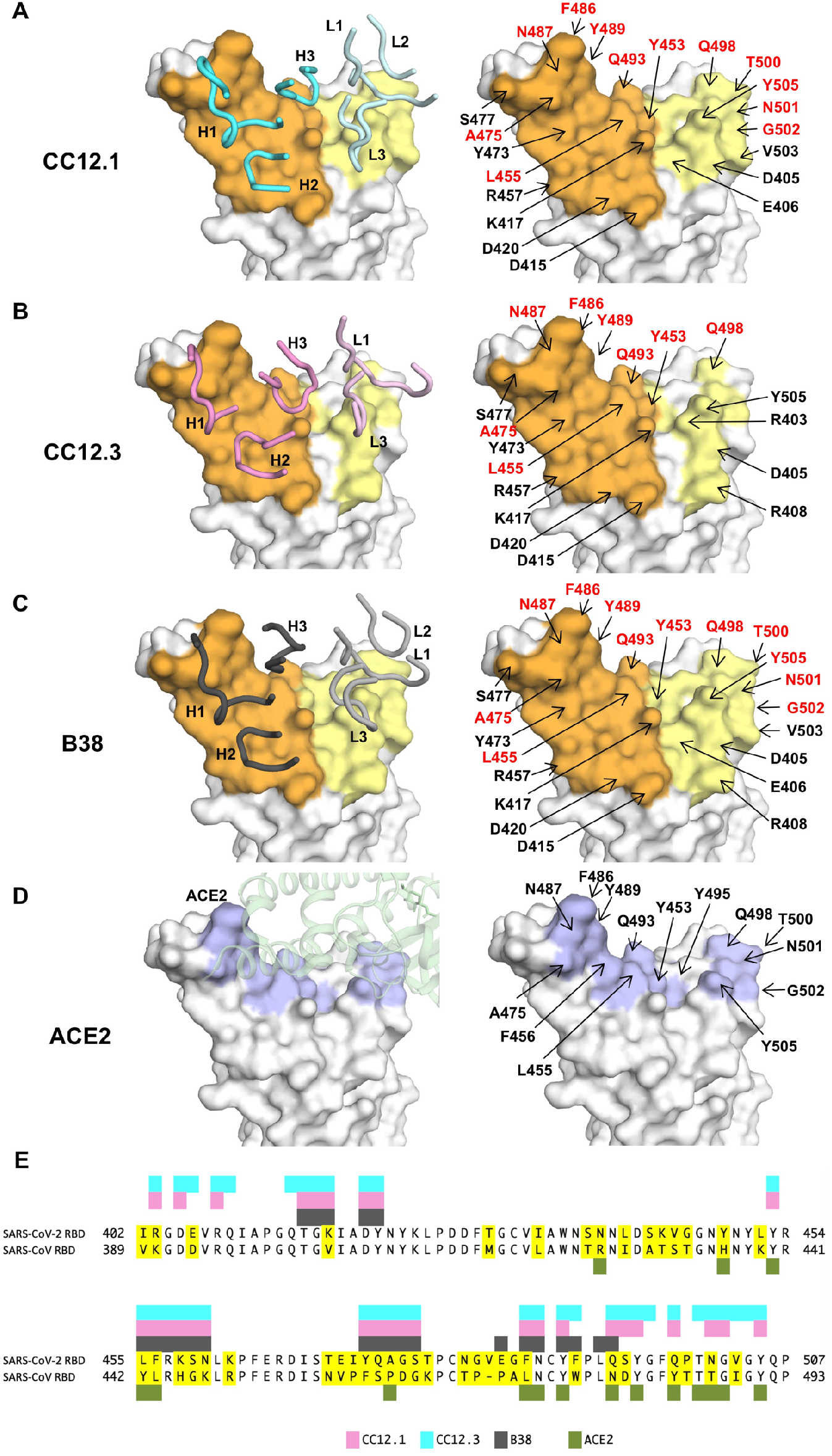
Epitopes of IGHV35-3 antibodies. **(A-C)** Epitopes of **(A)** CC12.1, **(B)** CC12.3, and **(C)** B38 (PDB 7BZ5) (*23*). Epitope residues contacting the heavy chain are in orange and the light chain are in yellow. On the left panels, CDR loops are labeled. On the right panels, epitope residues are labeled. For clarity, only representative epitope residues are labeled. Epitope residues that are also involved in ACE2 binding are in red. **(D)** ACE2-binding residues are shown in blue. On the left panel, ACE2 is shown in green with in semi-transparent representation. On the right panel, ACE2-binding residues are labeled. A total of 16 residues are used for ACE2 binding (*12*), but only 13 are labeled here since the other three are at the back of the structure in this view and do not interact with the antibodies of interest. **(E)** Epitope residues for CC12.1, CC12.3, and B38 were identified by PISA (*39*) and annotated on the SARS-CoV-2 RBD sequence, which is aligned to the SARS-CoV RBD sequence with non-conserved residues highlighted. The 16 ACE2-binding residues were as described previously (*12*).

To understand why IGHV3-53 is elicited as a public antibody response, the molecular interactions between the RBD and the heavy chains of CC12.1, CC12.3, and B38 were analyzed. The complementarity-determining regions (CDR) H1 and H2 of these antibodies interact extensively with the RBD mainly through specific hydrogen bonds (Fig. 3A-B). Interestingly, all residues on CDR H1 and H2 that hydrogen bond with the RBD are encoded by the germline IGHV3-53 (Fig. S1 and S7, Table S3). These interactions are almost identical among CC12.1, CC12.3, and B38 with the only difference at V_H_ residue 58. A somatic mutation V_H_ Y58F is present in both CC12.1 and CC12.3, but not in B38 (Fig. 3A-C, boxed residues). Nevertheless, this somatic mutation is unlikely to be essential for IGHV3-53 to engage the RBD, since V_H_ residue 58 in B38 still interacts with the RBD through an additional hydrogen bond (Fig. 3C). Of note, none of these antibody interactions mimic ACE2 binding (Fig. 3D).

**Figure 3.**
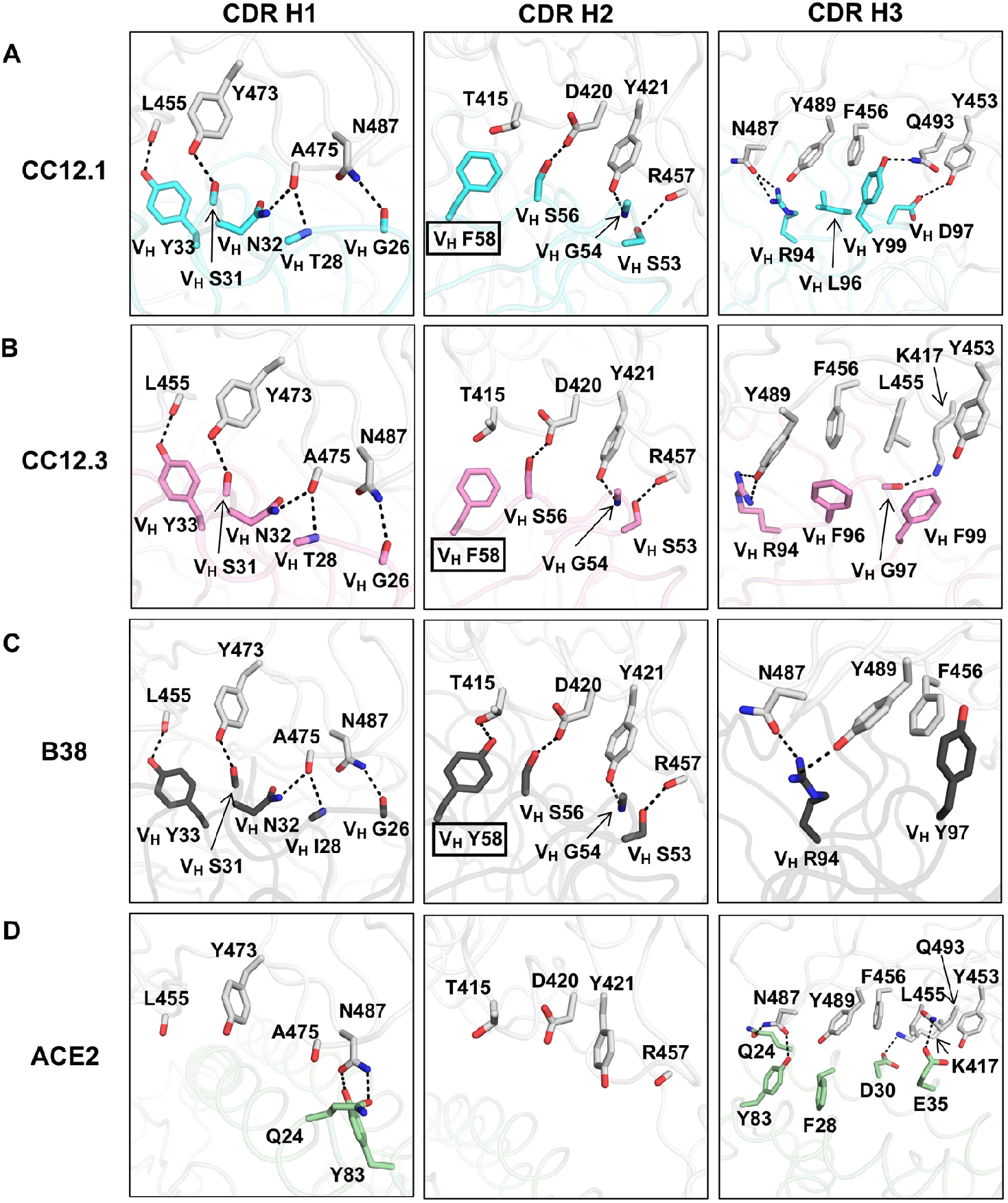
Interactions between the RBD and the heavy chain CDR loops. **(A-C)** Highly similar interaction modes between SARS-CoV-2 RBD and the antibody CDR H1 and H2 loops, but not the H3 loop are observed for in **(A)** CC12.1, **(B)** CC12.3, and **(C)** B38 (PDB 7BZ5) (*23*). The RBD is in white and antibody residues are in cyan, pink, and dark gray, respectively. Oxygen atoms are in red, and nitrogen atoms in blue. Hydrogen bonds are represented by dashed lines. **(D)** The interaction between ACE2 (green) and residues of the RBD (PDB 6M0J) (*12*) that are shown in **(A-C)**.

Our structural analysis reveals two key motifs in the IGHV3-53 germline sequence that are important for RBD binding, namely an NY motif at V_H_ residues 32 and 33 in the CDR H1, and an SGGS motif at V_H_ residues 53 to 56 in the CDR H2 (Fig. S8). The side chain of V_H_ N32 in the NY motif forms a hydrogen bond with the backbone carbonyl of A475 on the RBD, and this interaction is stabilized by an extensive network of hydrogen bonds with other antibody residues as well as a bound water molecule (Fig. 4A). V_H_ N32 also hydrogen bonds with V_H_ R94, which in turn hydrogen bonds with N487 and Y489 on the RBD (Fig. 4A). These interactions enhance not only RBD-Fab interaction, but also stabilize CDR and framework residues and conformations. V_H_ Y33 in the NY motif inserts into a hydrophobic cage formed by RBD residues Y421, F456, L455 and the aliphatic component of K417 (Fig. 4B). A hydrogen bond between V_H_ Y33 and the carbonyl oxygen of L455 on RBD further strengthens the interaction. The second key motif SGGS in CDR H2 forms extensive hydrogen bond network with the RBD (Fig. 4C), including four hydrogen bonds that involve the hydroxyl side chains of V_H_ S53 and V_H_ S56, and four water-mediated hydrogen bonds to the backbone carbonyl of V_H_ G54, the backbone amide of V_H_ S56, and the side chain of V_H_ S56. Along with V_H_ Y52, the SGGS motif takes part in a type I beta turn, with a positive Φ-angle for V_H_ G55 at the end of the turn. In addition, the Cα of V_H_ G54 is only 4 Å away from the RBD, indicating that side chains of other amino acids would clash with the RBD if they were present at this position. As a result, the SGGS motif is a perfect fit for interacting with the RBD at this location.

**Figure 4.**
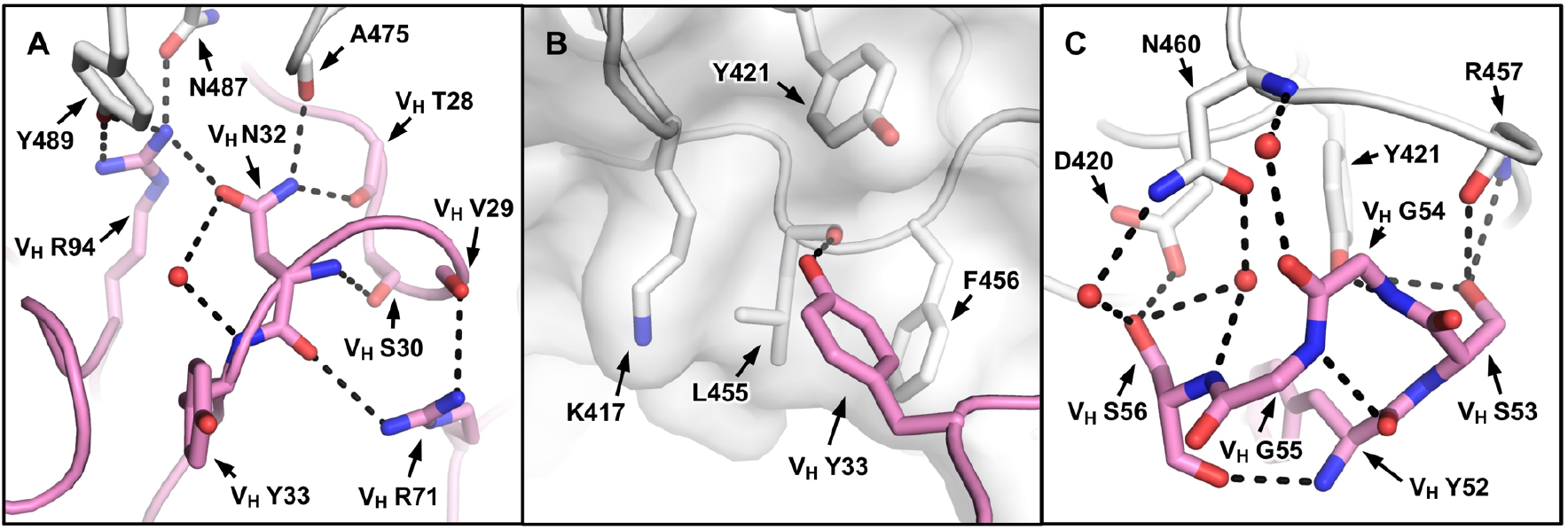
Two IGHV3-53 germline-encoded motifs. **(A)** The extensive hydrogen bond network that involves V_H_ N32 of the NY motif in CDR H1 is illustrated. **(B)** The hydrophobic cage interaction between the RBD and V_H_ Y33 of the NY motif in CDR H1 is shown. **(C)** The hydrogen bond network that involves the SGGS motif in CDR H2 is highlighted. CC12.3 is shown because its structure is at higher resolution than CC12.1.

Overall, these observations demonstrate the importance of the NY and SGGS motifs, which are both encoded in the IGHV3-53 germline, for engaging the RBD. In fact, besides IGHV3-53, the only other IGHV gene that contains an NY motif in CDR H1 and an SGGS motif in CDR H2 is IGHV3-66, which is a closely-related IGHV gene to IGHV3-53 (*32*). As compared to IGHV3-53, IGHV3-66 has a lower occurrence frequency in the repertoire of healthy individuals (0.3% to 1.7%) (*29*), which may explain why IGHV3-66 is less prevalent than IGHV3-53, but yet is still quite commonly observed (*19–22, 24, 26*) in antibodies in SARS-CoV-2 patients (Fig. 1A). Overall, our structural analysis has identified two germline-encoded binding motifs that enable IGHV3-53 to act as a public antibody and target the SARS CoV-2 RBD with no mutations required from affinity maturation.

While the binding mode of CDR H1 and H2 to RBD is highly similar among CC12.1, CC12.3, and B38, CDR H3 interaction with the RBD is different (Fig. 3A-C) due to differences in the CDR H3 sequences and conformations (Fig. 5A-B). For example, while CDR H3 of CC12.1 can interact with RBD Y453 through a hydrogen bond, CDR H3s of CC12.3 and B38 do not from such a hydrogen bond (Fig. 3A-C). Similarly, due to the difference in light-chain gene usage, the light-chain interactions with the RBD can vary substantially in IGHV3-53 antibodies (Fig. S9). Overall, our structural analysis demonstrates that IGHV3-53 provides a highly versatile framework for antibodies to target the ACE2 binding site in SARS-CoV-2 RBD.

**Figure 5.**
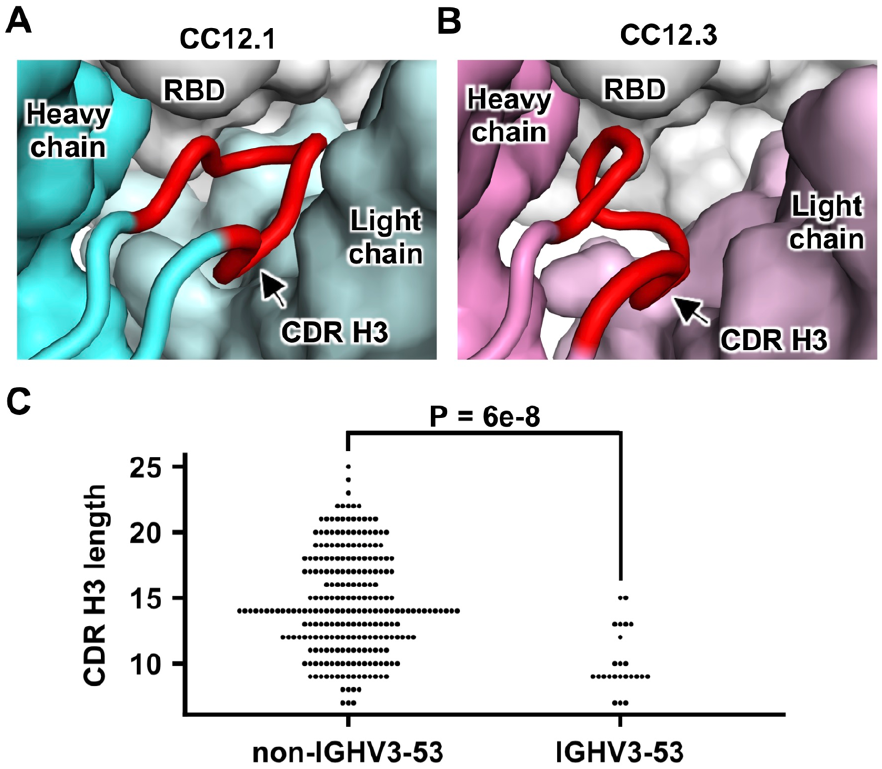
Constraints on CDR H3 length. **(A)** The heavy and light chains of CC12.1 (cyan), as well as the RBD (white) are shown in surface representation, with CDR H3 (red) highlighted in cartoon representation. **(B)** Same as panel A, except that CC12.3 (pink) is shown. **(C)** The lengths of CDR H3 in RBD-targeting antibodies that were previously isolated (*17–28*) are analyzed. The distribution of CDR H3 lengths in RBD-targeting IGHV3-53 antibodies and those in non-IGHV3-53-encoded antibodies are compared. A Mann-Whitney U test was performed to compute the p-value.

An interesting feature of CC12.1 and CC12.3 is their relatively short CDR H3. While the CDR H3 sequences of CC12.1 and CC12.3 are very different, both have a length of nine amino acids (Kabat numbering), whereas the average CDR H3 length for human antibodies is around 13 (*33*), as compared to for example very long CDR H3s (up to 30 residues) on average that seem to be required for many broadly neutralizing antibodies to HIV-1 (*34*). Similarly, antibody B38 has a very short CDR H3 length of seven residues (*23*). It is unlikely that longer CDR H3’s can be accommodated in these antibodies since their epitopes are relatively flat with no large pocket to insert a protruding CDR (Fig. 2A-C). This conclusion was also arrived in a recent study that also reported SARS-CoV-2 RBD-targeting antibodies that are encoded by either IGHV3-53 or IGHV3-66 tend to have a short CDR H3 (*28*). In fact, IGHV3-53 and IGHV3-66 antibodies in general have slightly shorter than average CDR H3’s (by around one residue), but do appear to have a few much shorter CDR H3’s (<10 amino acids) than average in the baseline antibody repertoire (*30*). In CC12.1 and CC12.3, the space for CDR H3 to fit in the interface within the RBD is limited (Fig. 5A-B), which thus constrains its length. Consistently, among RBD-targeting antibodies currently reported (*17–28*), those encoded by IGHV3-53 have a significantly shorter CDR H3 compared to those encoded by other IGHV genes (p-value = 6e-8, Mann-Whitney U test) (Fig. 5C). These observations provide structural and statistical evidence that a short CDR H3 length is a molecular feature of the IGHV3-53-encoded public antibody response to the SARS-CoV-2 RBD, reminiscent of a short 5-residue CDR L3 in IGHV1-2 antibodies to the CD4 receptor binding site in gp120 of HIV-1 Env (*35*).

Besides IGHV3-53, several other IGHV genes such as IGHV1-2, IGHV3-9, and IGHV3-30 are also more frequently observed than other germlines in SARS-CoV-2 RBD-targeting antibodies (Fig. 1A). The molecular mechanisms of these antibody responses to SARS-CoV-2 will need to be characterized in the future. In addition, whether other antibody germline gene segments, including the IGHD and the light chain, contribute to public antibody responses to SARS-CoV-2 will also need to be further addressed. Notwithstanding, the detailed characterization of this public antibody response to SARS CoV-2 is already be a promising starting point for rational vaccine design (*36*), especially given limited to no affinity maturation is required from the germline to achieve a high affinity neutralizing antibody response to the RBD. In addition, IGHV3-53 exists at a reasonable frequency in healthy individuals (*29, 30*), indicating that this public antibody could be commonly elicited during vaccination (*37*), and aid in design of both antibody and small molecule therapeutics (*7, 38*).

## ACKNOWLEDGEMENTS

We thank Robyn Stanfield for assistance in data collection and Bryan Briney for naïve antibody germline analysis. We are grateful to the staff of Stanford Synchrotron Radiation Laboratory (SSRL) Beamline 12-1 for assistance. This work was supported by NIH K99 AI139445 (N.C.W.), the Bill and Melinda Gates Foundation OPP1170236 (I.A.W. and D.R.B.), NIH CHAVD (UM1 AI44462 to I.A.W., D.S. and D.R.B.), and the IAVI Neutralizing Antibody Center. Use of the SSRL, SLAC National Accelerator Laboratory, is supported by the U.S. Department of Energy, Office of Science, Office of Basic Energy Sciences under Contract No. DE-AC02-76SF00515. The SSRL Structural Molecular Biology Program is supported by the DOE Office of Biological and Environmental Research, and by the National Institutes of Health, National Institute of General Medical Sciences (including P41GM103393).

## AUTHOR CONTRIBUTIONS

M. Y., H.L., N.C.W., F.Z., D.H., T.F.R., E.L., D.S, J.G.J., D.R.B. and I.A.W. conceived and designed the study. M.Y., H.L., N.C.W., C.C.D.L., W.Y. and Y.H. expressed and purified the proteins. M.Y. and C.C.D.L. performed biolayer interferometry binding assays. M.Y., H.L., N.C.W., X.Z. and H.T. performed the crystallization and X-ray data collection. M.Y. and X.Z. determined and refined the X-ray structures. M.Y., H.L., N. C.W., C.C.D.L. and X.Z. analyzed the data. M.Y., H.L., N.C.W. and I.A.W. wrote the paper and all authors reviewed and/or edited the paper.

## MATERIALS AND METHODS

### Expression and purification of SARS-CoV-2 RBD

The receptor-binding domain (RBD) (residues 319-541) of the SARS-CoV-2 spike (S) protein (GenBank: QHD43416.1) was cloned into a customized pFastBac vector (*40*), and fused with an N-terminal gp67 signal peptide and C-terminal His6 tag (*17*). A recombinant bacmid DNA was generated using the Bac-to-Bac system (Life Technologies). Baculovirus was generated by transfecting purified bacmid DNA into Sf9 cells using FuGENE HD (Promega), and subsequently used to infect suspension cultures of High Five cells (Life Technologies) at an MOI of 5 to 10. Infected High Five cells were incubated at 28 °C with shaking at 110 r.p.m. for 72 h for protein expression. The supernatant was then concentrated using a 10 kDa MW cutoff Centramate cassette (Pall Corporation). The RBD protein was purified by Ni-NTA, followed by size exclusion chromatography, and buffer exchanged into 20 mM Tris-HCl pH 7.4 and 150 mM NaCl.

### Expression and purification of Fabs

For CC12.1 and CC12.3, the heavy and light chains were cloned into phCMV3. The plasmids were transiently co-transfected into ExpiCHO cells at a ratio of 2:1 (HC:LC) using ExpiFectamine™ CHO Reagent (Thermo Fisher Scientific) according to the manufacturer’s instructions. The supernatant was collected at 10 days post-transfection. The Fabs were purified with a CaptureSelect™ CH1-XL Affinity Matrix (Thermo Fisher Scientific) followed by size exclusion chromatography. CR3022 was expressed and purified as described previously (*17*).

### Expression and purification of ACE2

The N-terminal peptidase domain of human ACE2 (residues 19 to 615, GenBank: BAB40370.1) was cloned into phCMV3 vector, and fused with a C-terminal Fc tag. The plasmids were transiently transfected into Expi293F cells using ExpiFectamine™ 293 Reagent (Thermo Fisher Scientific) according to the manufacturer’s instructions. The supernatant was collected at 7 days post-transfection. Fc-tagged ACE2 protein was then purified with a Protein A column (GE Healthcare) followed by size exclusion chromatography.

### Crystallization and structural determination

CC12.1/RBD, CC12.3/RBD, CC12.1/CR3022/RBD, and CC12.3/CR3022/RBD complexes were formed by mixing each of the protein components at an equimolar ratio and incubated overnight at 4°C. Each complex was adjusted to 13 mg/ml and screened for crystallization using the 384 conditions of the JCSG Core Suite (Qiagen) and ProPlex screen (Molecular Dimensions) on either our custom-designed robotic CrystalMation system (Rigaku) or an Oryx8 (Douglas Instruments) at Scripps Research. Crystallization trials were set-up by the vapor diffusion method in sitting drops containing 0.1 μl of protein and 0.1 μl of reservoir solution. Diffraction-quality crystals were obtained in the following conditions:

CC12.1/RBD complex (13 mg/ml): 0.1M sodium citrate pH 5.5 and 15% (w/v) polyethylene glycol 6000 at 20°C
CC12.3/RBD complex (13 mg/mL): 0.1 M sodium phosphate pH 6.5 and 12% (w/v) polyethylene glycol 8000 at 20°C
CC12.1/RBD/CR3022 complex (13 mg/mL): 20% PEG-3000, 0.2 M sodium chloride, 0.1 M HEPES pH 7.5 at 20°C
CC12.3/RBD/CR3022 complex (13 mg/mL): 0.1M Tris pH 8, 15% ethylene glycol, 1M lithium chloride, 10% PEG 6000 at 20°C

Of note, these four complexes crystallized in a broad range of pHs. All crystals appeared on day 3 and were harvested on day 7. Before flash cooling in liquid nitrogen for X-ray diffraction studies, crystals were equilibrated in reservoir solution supplemented the following cryoprotectants:

CC12.1/RBD complex: 20% glycerol
CC12.3/RBD complex: 20% glycerol
CC12.1/RBD/CR3022 complex: 10% ethylene glycol
CC12.3/RBD/CR3022 complex: none were required

Diffraction data were collected at cryogenic temperature (100 K) at Stanford Synchrotron Radiation Lightsource (SSRL) on the new Scripps/Stanford beamline 12-1 with a beam wavelength of 0.97946 Å, and processed with HKL2000 (*41*). Structures were solved by molecular replacement using PHASER (*42*) with PDB 6YLA (*43*), 4TSA, and 4ZD3 (*44*). Iterative model building and refinement were carried out in COOT (*45*) and PHENIX (*46*), respectively. Epitope and paratope residues, as well as their interactions, were identified by accessing PISA at the European Bioinformatics Institute (http://www.ebi.ac.uk/pdbe/prot_int/pistart.html) (*39*).

### Biolayer interferometry binding assay

Antibody binding and competition assays were performed by biolayer interferometry (BLI) using an Octet Red instrument (FortéBio) as described previously (*47*), with Ni-NTA biosensors. There were five steps in the assay: 1) baseline: 60s with 1x kinetics buffer; 2) loading: 180 s with 20 μg/mL of 6x His-tagged SARS-CoV-2 RBD proteins; 3) baseline: 135 s with 1x kinetics buffer; 4) association: 240 s with serial diluted concentrations of CC12.1 Fab or CC12.3 Fab; and 5) dissociation: 240 s with 1x kinetics buffer. For K_d_ estimation, a 1:1 binding model was used.

For competition assays, CC12.1 Fab, CC12.3 Fab, CR3022 Fab, and human ACE2-Fc were all diluted to 250 nM. Ni-NTA biosensors were used. In brief, the assay has five steps: 1) baseline: 60s with 1x kinetics buffer; 2) loading: 120 s with 20 μg/mL, 6x His-tagged SARS-CoV-2 RBD proteins; 3) baseline: 120 s with 1x kinetics buffer; 4) first association: 180 s with CC12.1 Fab, CC12.3 Fab or CR3022 Fab; and 5) second association: 180 s with CC12.1 Fab, CC12.3 Fab, CR3022 Fab, or human ACE2-Fc or disassociation with 1x kinetics buffer for each first association.

---

**Fig. S1.**
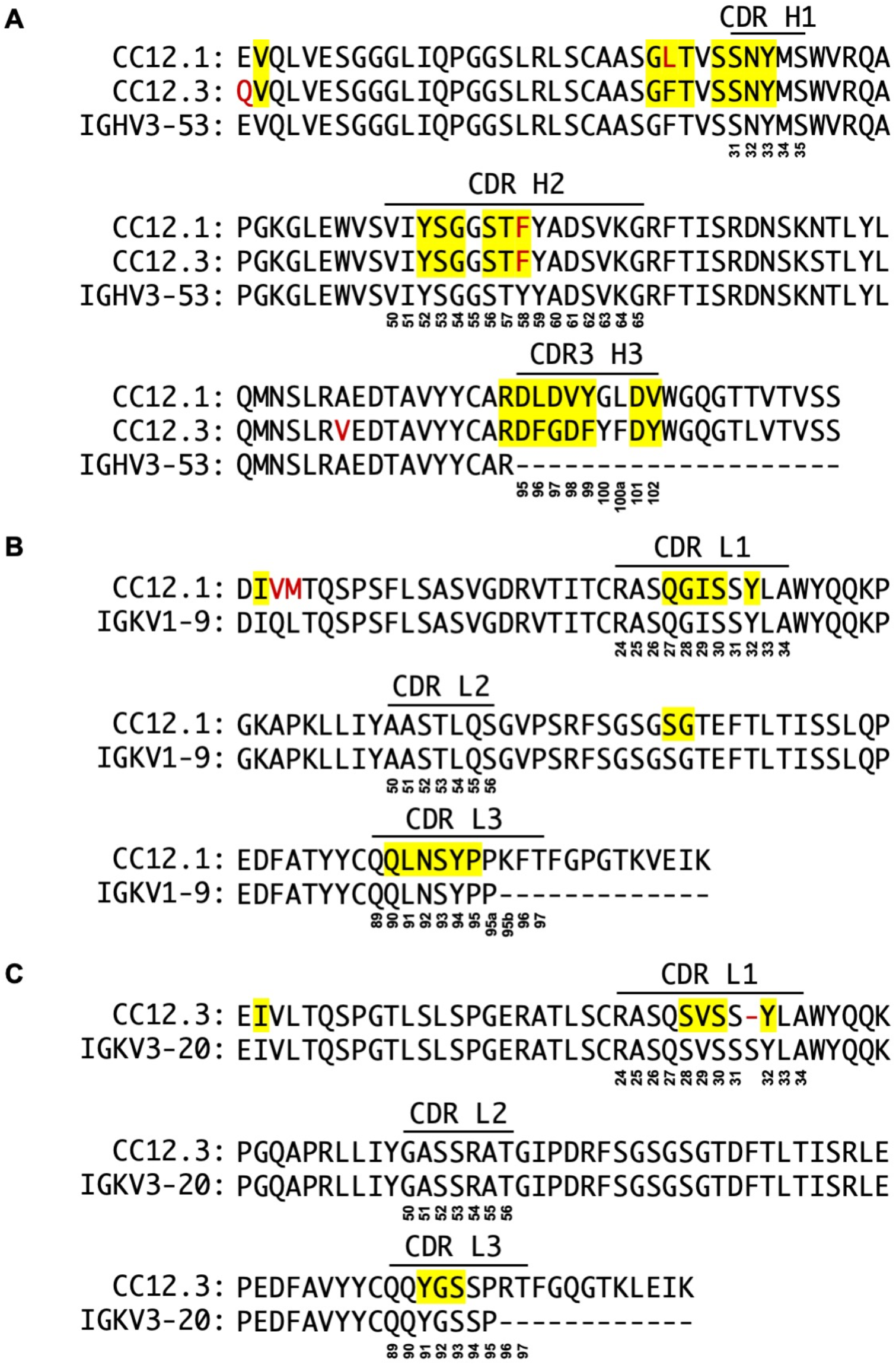
Comparison of CC12.1 and CC12.3 sequence to the IGHV3-53 germline sequence. **(A)** Alignment of the heavy chain variable domain sequences of CC12.1 and CC12.3 with the germline IGHV3-53 sequence **(B)** Alignment of the light-chain variable domain sequence of CC12.1 with the germline IGKV1-9 sequence. **(C)** Alignment of the light-chain variable domain sequence of CC12.3 with the germline IGKV3-20 sequence. The regions that correspond to CDR H1, H2, H3, L1, L2, and L3 are indicated. Residues that differ from the germline are highlighted in red. Residue positions in the CDRs are labeled according to the Kabat numbering scheme. Residues that interact with the RBD are highlighted in yellow.

**Fig. S2.**
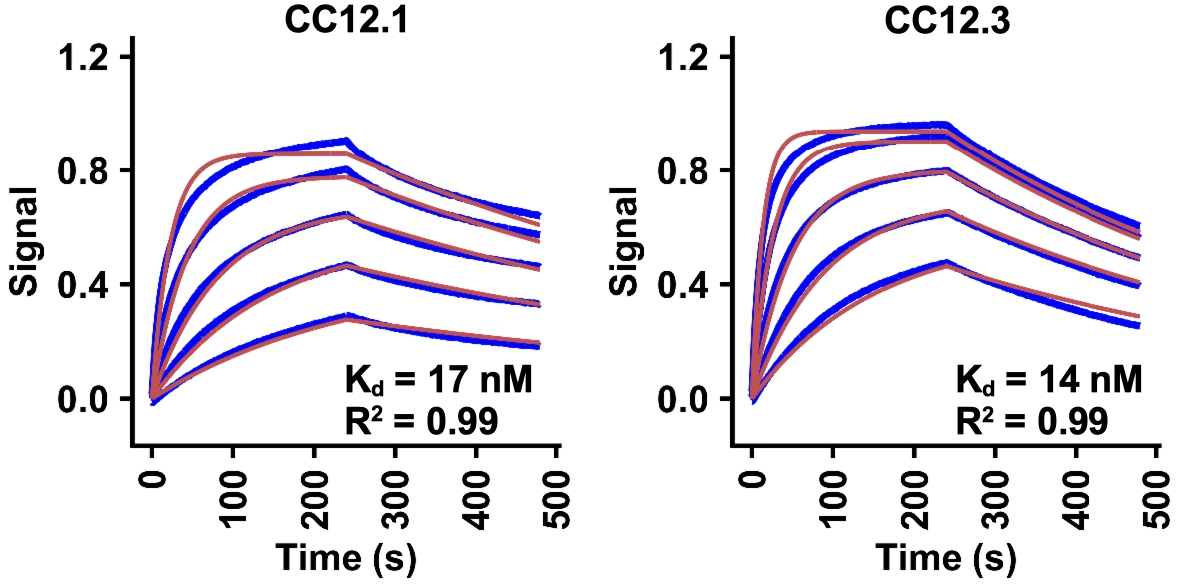
Sensorgrams for binding of CC12.1 and CC12.3 Fabs to SARS-CoV-2 RBD. Binding kinetics of CC12.1 and CC12.3 Fab against SARS-CoV-2 RBD were measured by biolayer interferometry (BLI). Y-axis represents the response. Blue lines represent the response curves and red lines represent the 1:1 binding model. Binding kinetics were measured for five concentrations of Fab at 2-fold dilution ranging from 500 nM to 31.25 nM. The K_d_ and R^2^ of the fitting are indicated.

**Fig. S3.**
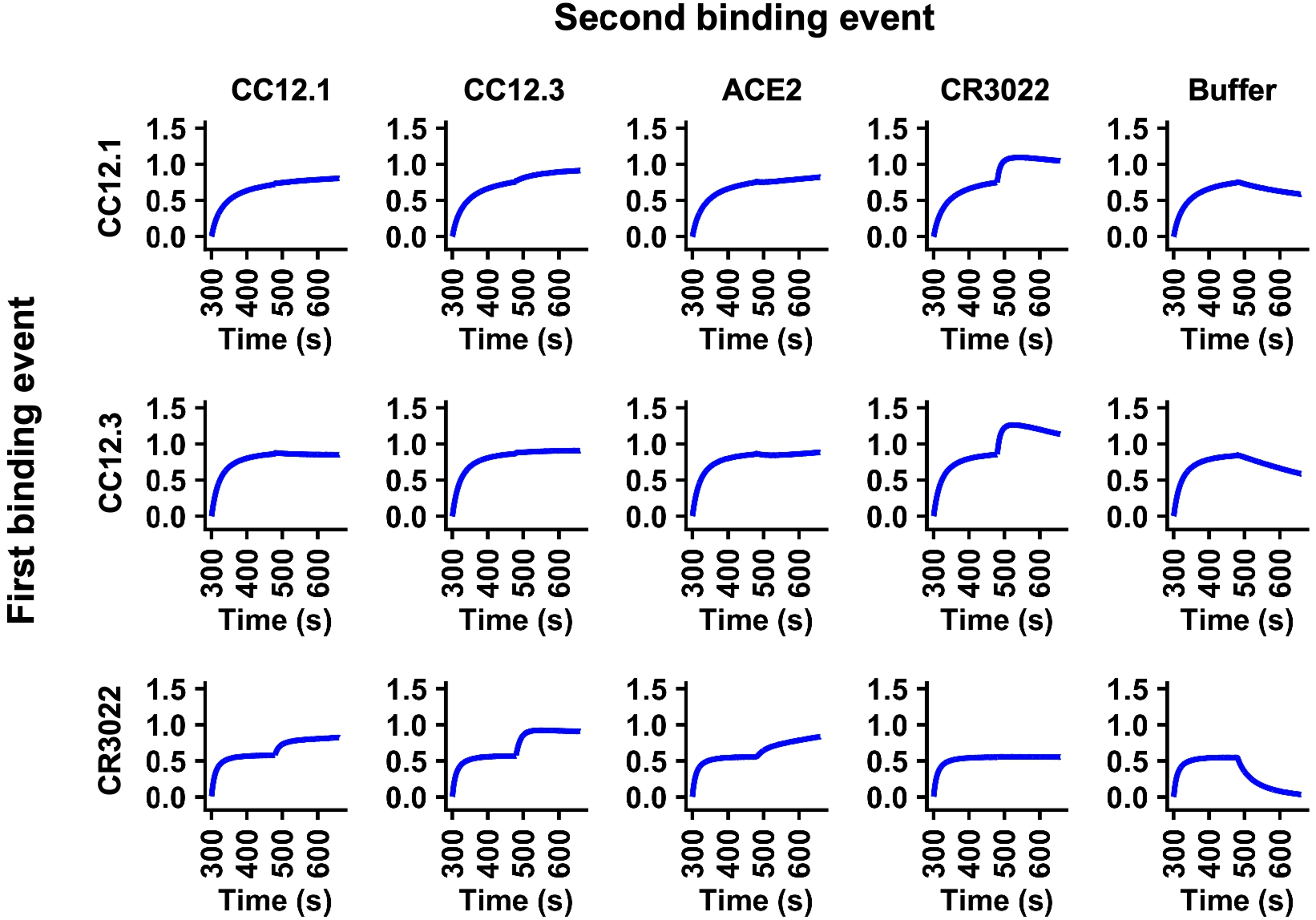
Competition assay between different Fabs and ACE2. Competition between CC12.1, CC12.3, CR3022, and ACE2 was measured by biolayer interferometry (BLI). Y-axis represents the response. The biosensor was first loaded with SARS-CoV-2 RBD, followed by two binding events: 1) CC12.1, CC12.3, or CR3022, and 2) CC12.1, CC12.3, or CR3022, ACE2, and buffer (negative control). A period of 180 s was used for each of the binding events. A further increase in signal during the second binding event (starting at 480 s time point) indicates lack of competition with the first ligand.

**Fig. S4.**
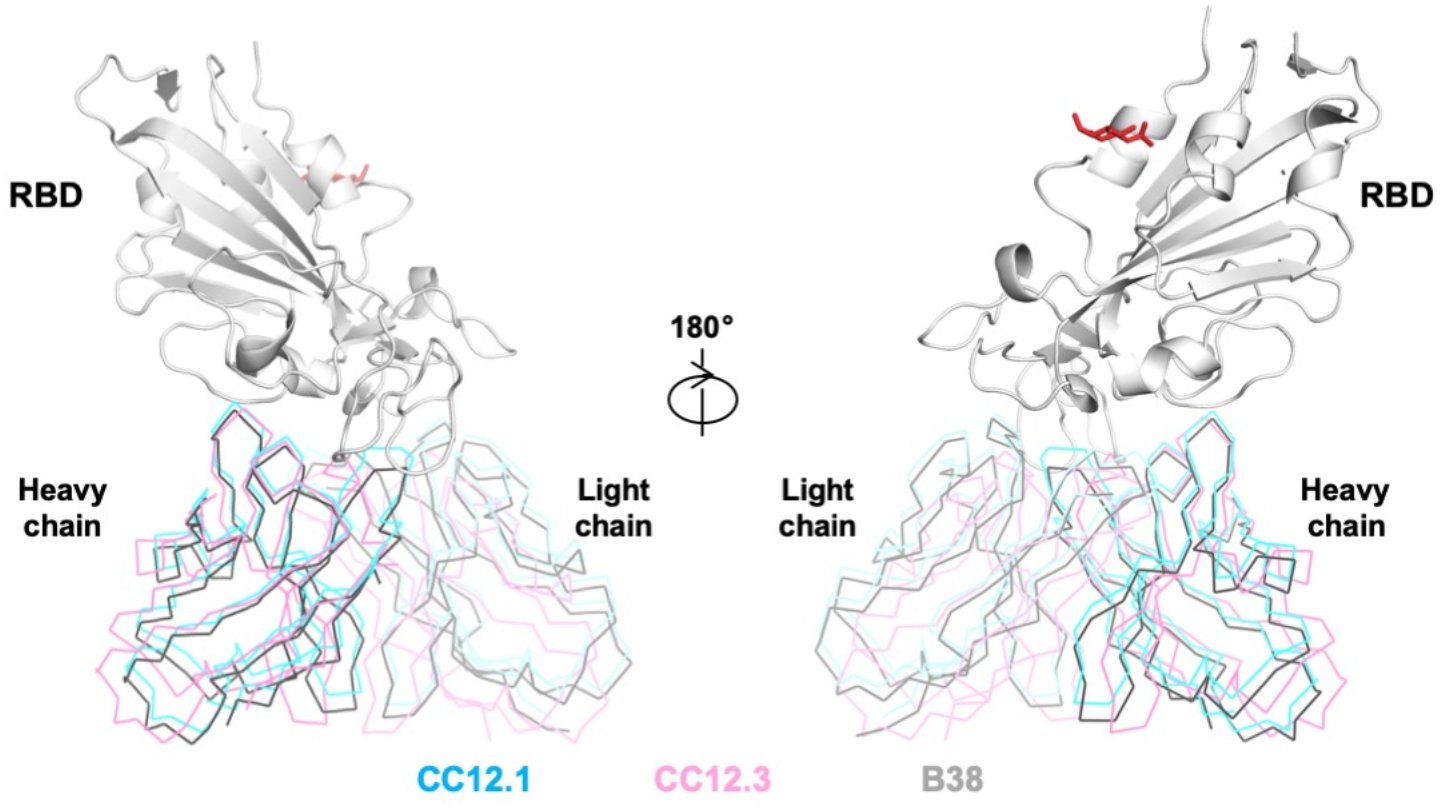
Structural comparison of the binding modes among IGHV3-53 antibodies. The binding modes of CC12.1 (cyan), CC12.3 (pink), and B38 (gray) to SARS-CoV-2 (white) are compared. B38 in complex with SARS-CoV-2 RBD was from PDB 7BZ5 (*23*). The N-glycan observed at SARS-CoV-2 RBD N343 is shown in red.

**Fig. S5.**
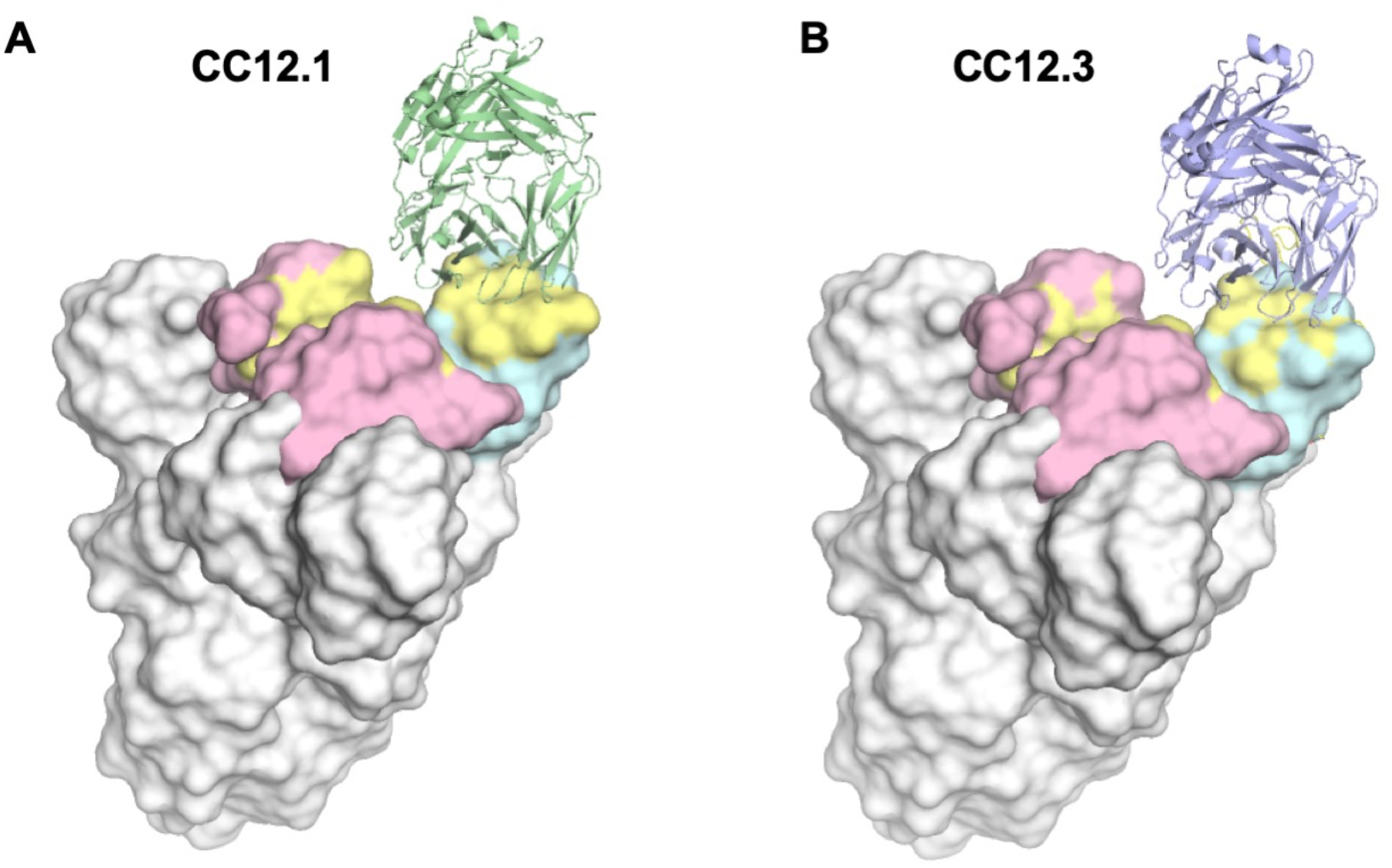
Modelling the binding of CC12.1 and CC12.3 on the homotrimeric spike (S) protein. The S trimer is shown with one RBD in the up conformation (cyan) and two RBDs in the down conformation (pink). The CC12.1 and CC12.3 epitopes are shown in yellow. **(A)** Model of the binding of CC12.1 (green) to the RBD up conformation. **(B)** Model of the binding of CC12.3 (blue). PDB 6VSB is used in the modeling (*48*). The complete epitopes of CC12.1 and CC12.3 are accessible only when the RBD is in the up, but not the down, conformation.

**Fig. S6.**
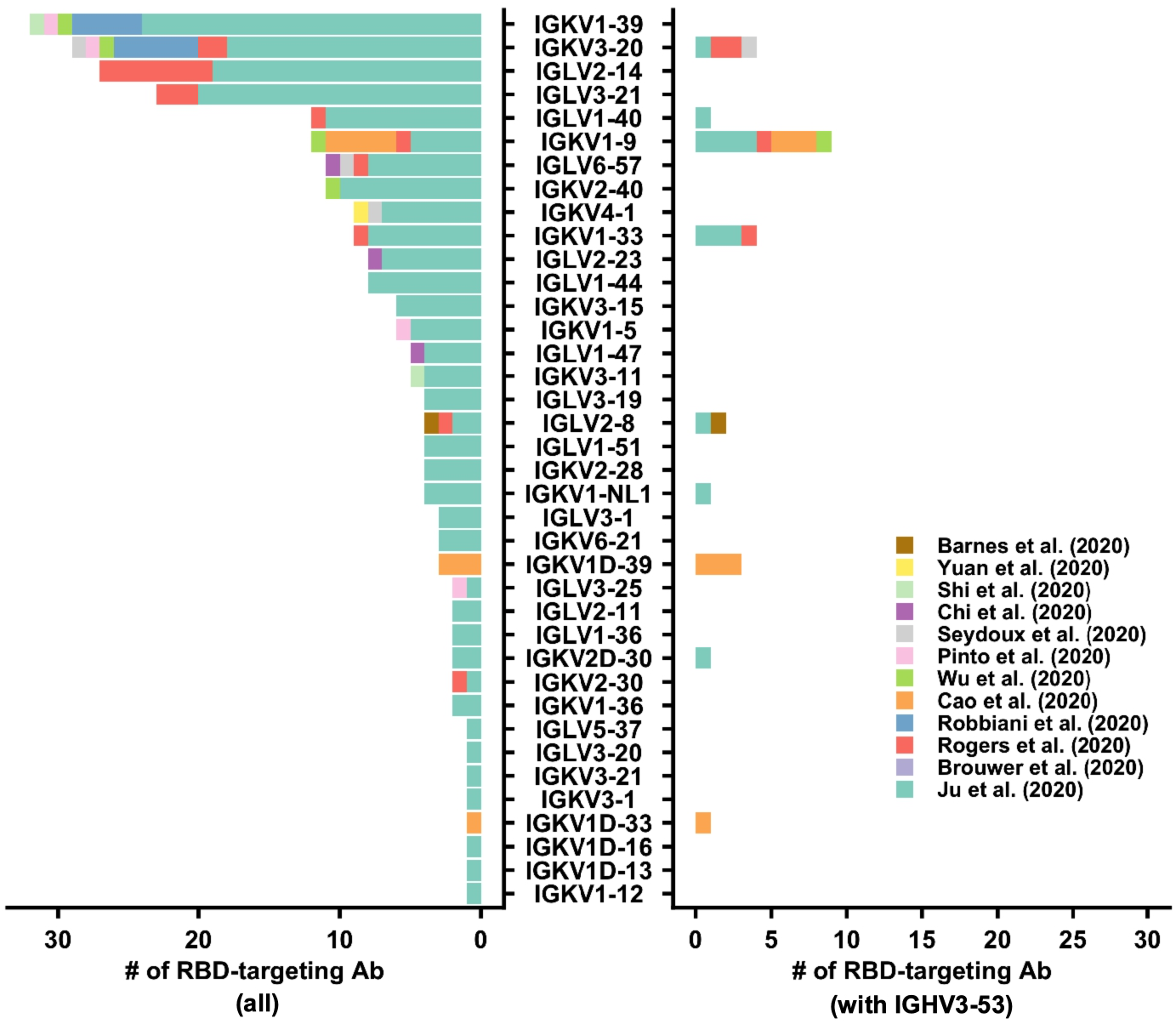
Light-chain germline gene use in SARS-CoV-2 RBD-targeting antibodies. The distribution of light-chain germline gene use of SARS-CoV-2 RBD-targeting antibodies that have been recently isolated (*17–28*) is shown on the left. The distribution of light-chain germline gene use in antibodies that target SARS-CoV-2 RBD to the subset of antibodies that pair with IGHV3-53 is shown on the right.

**Fig. S7.**
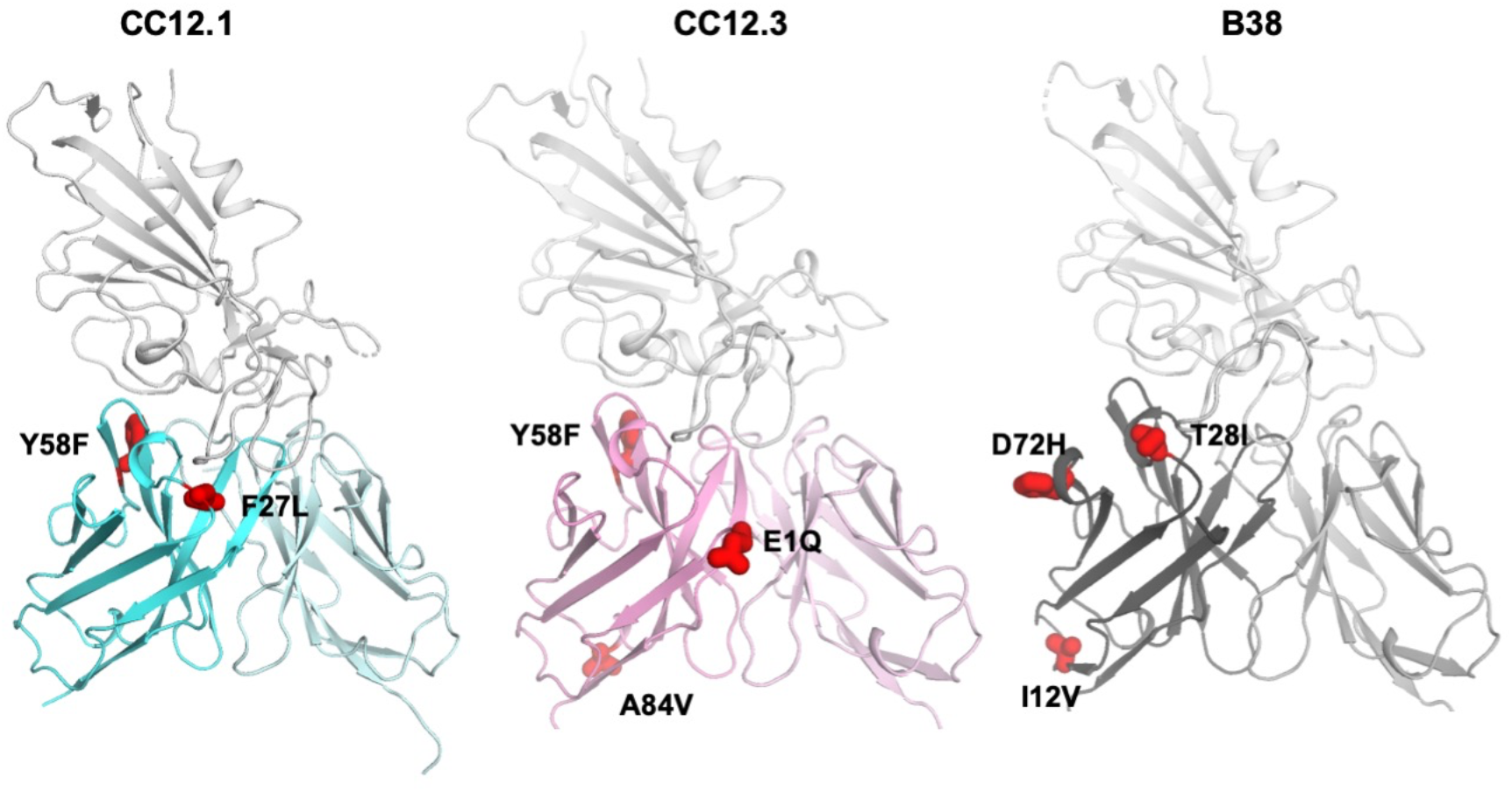
Locations of heavy chain somatic mutations. Somatic mutations on the heavy chains of CC12.1, CC12.3, and B38 are labeled and shown in red on the structure. Somatic mutations contribute minimally to the antibody binding interactions.

**Fig. S8.**
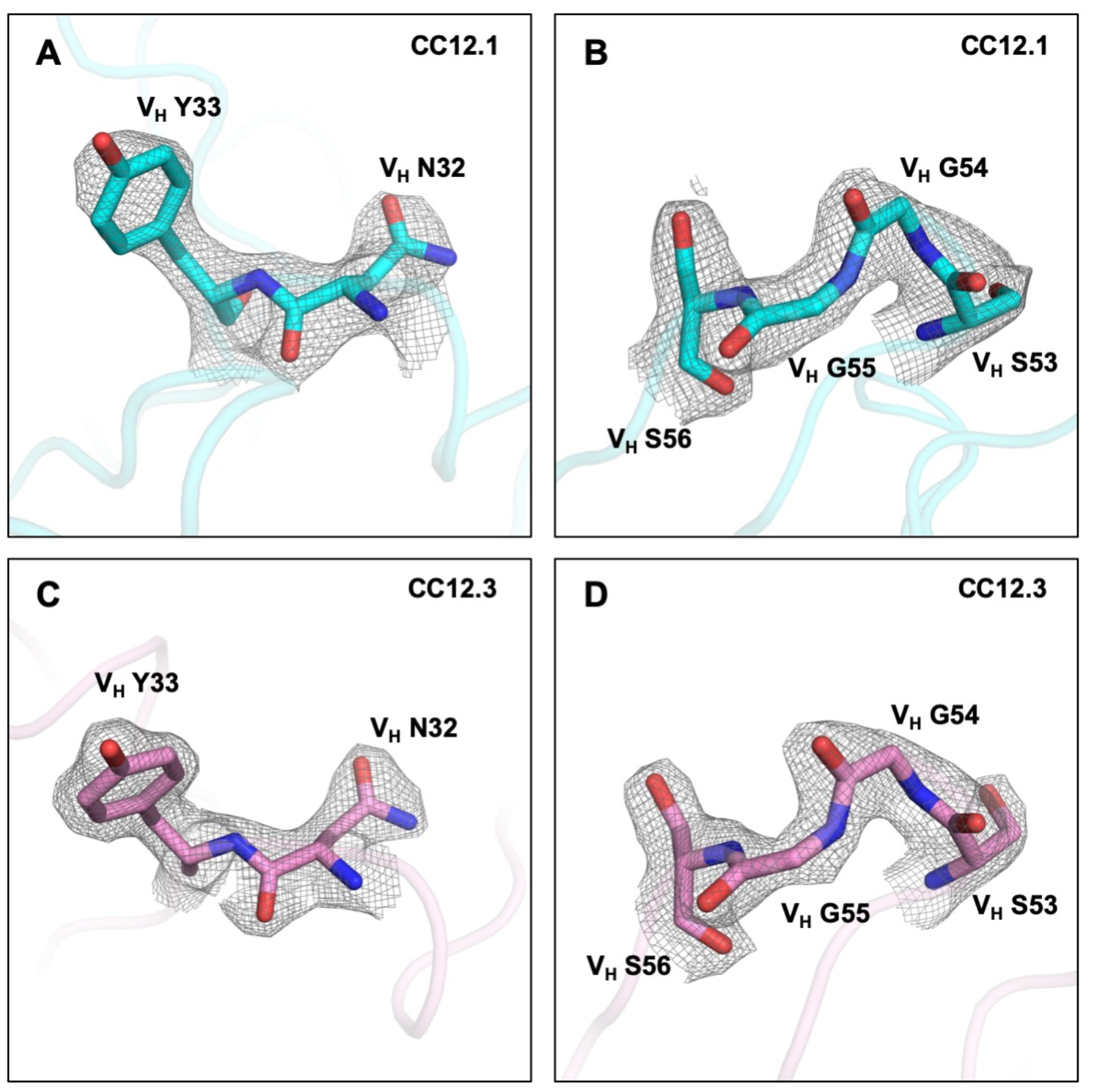
Electron density maps for IGHV3-53-encoded paratope regions of CC12.1 and CC12.3. **(A-B)** Final 2Fo-Fc electron density maps for the IGHV3-53-encoded paratope regions around V_H_ N32 and Y33 (CDR H1) and V_H_ S53 to S56 (CDR H2) of CC12.1, both contoured at 1.2 σ. **(C-D)** Final 2Fo-Fc electron density maps for IGHV3-53-encoded paratope regions around VH N32 and Y33 and VH S53 to S56 of CC12.3, both contoured at 1.8 σ.

**Fig. S9.**
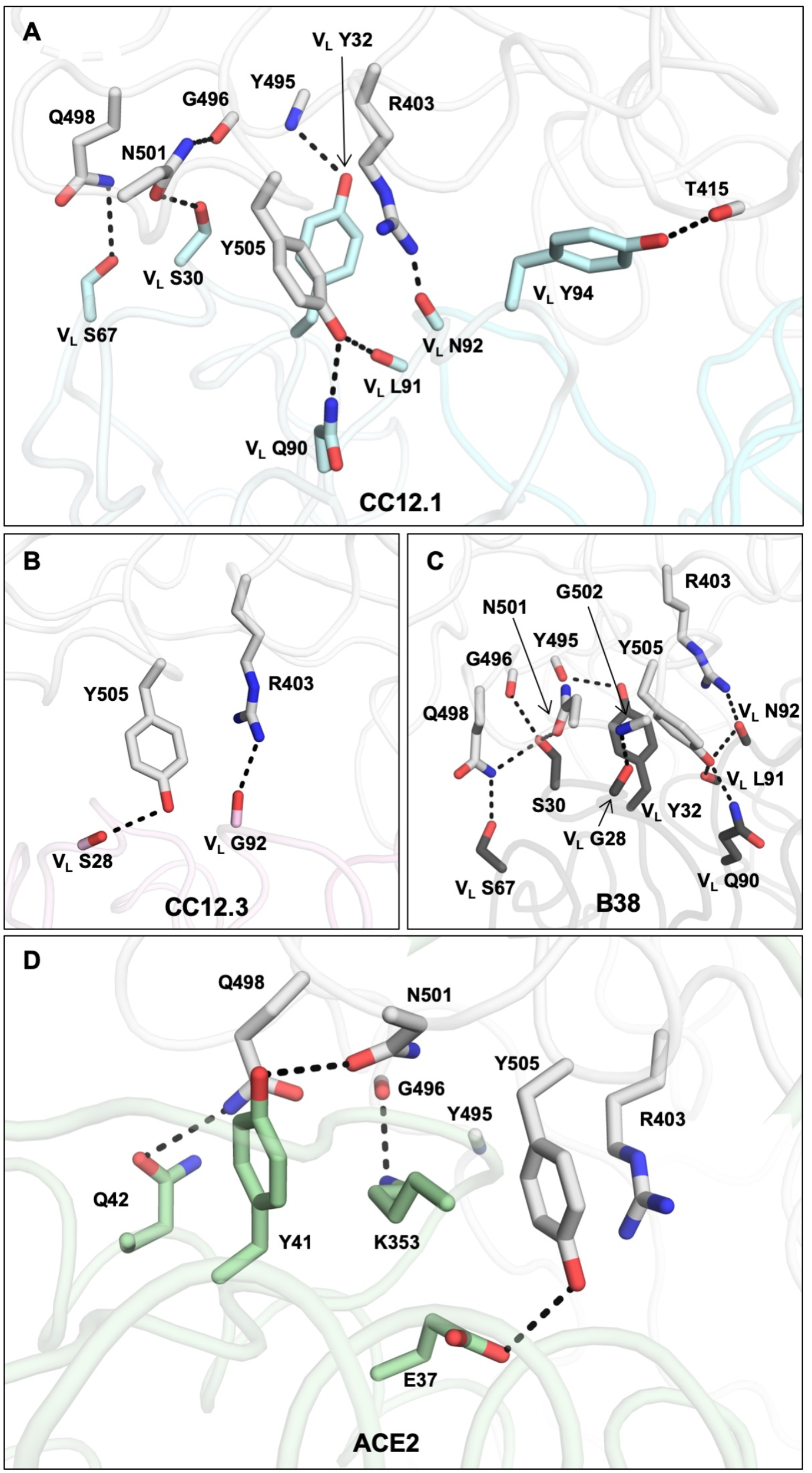
Interactions between the light chain and the RBD. **(A-C)** Representative interactions between SARS-CoV-2 RBD and the light chain in **(A)** CC12.1, **(B)** CC12.3, and **(C)** B38 (PDB 7BZ5) (*23*) are shown. RBD is in white. Oxygen atoms are in red. Nitrogen atoms are in blue. Hydrogen bonds are represented by dashed lines. The light chains from both CC12.1 and B38 form an extensive hydrogen bond network with the RBD, whereas the interaction between the light chain of CC12.3 and the RBD is minimal. **(D)** The interaction between ACE2 and RBD residues (PDB 6M0J) (*12*) that are shown in **(A-C)**. None of the interactions between the light chain and RBD mimic those between ACE2 and RBD.

**Table S1.**
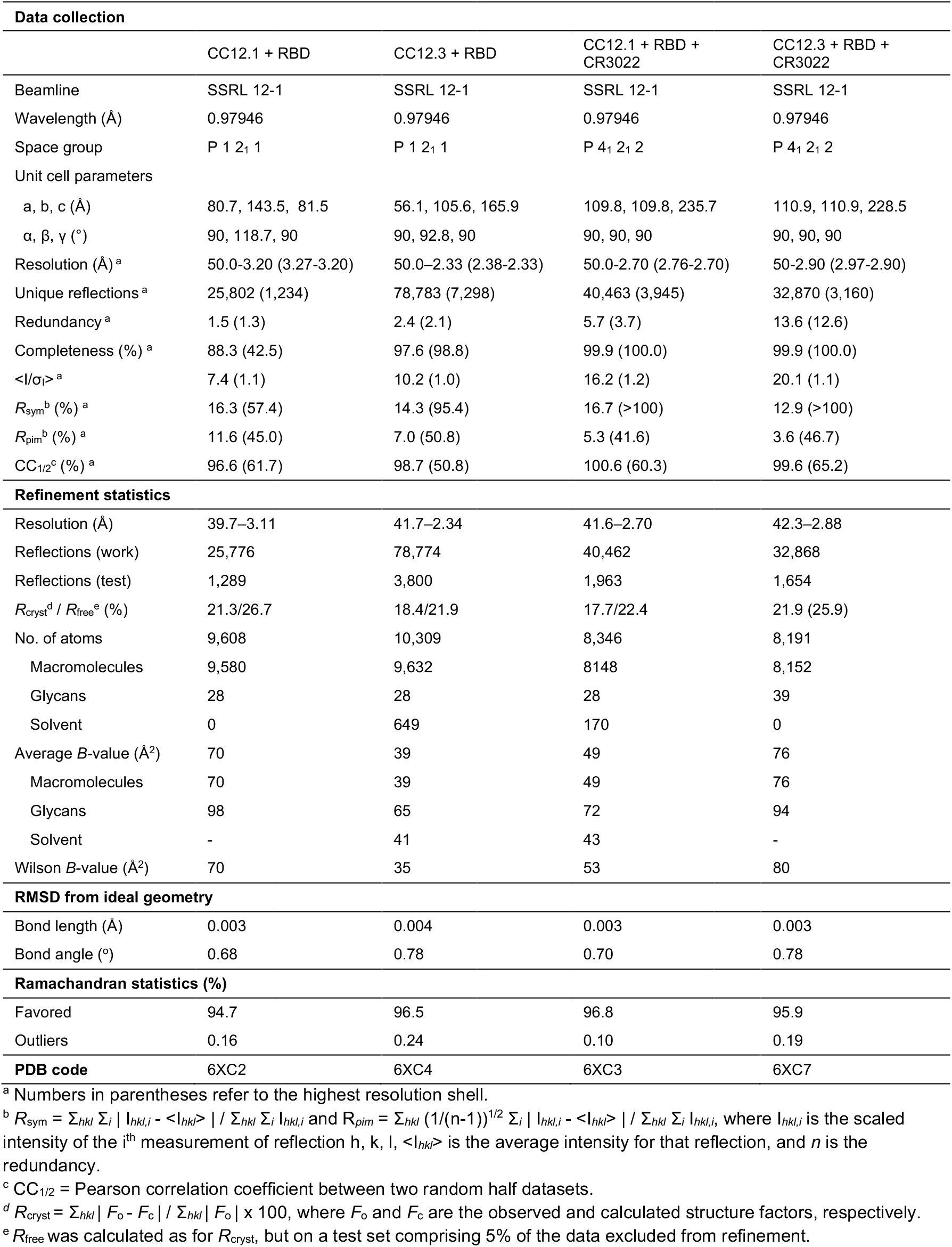
X-ray data collection and refinement statistics

**Table S2.** A list of previously reported SARS-CoV-2 RBD-targeting antibodies.

| Antibody name | Heavy chain | Light chain | CDR H3 length | Reference |
| --- | --- | --- | --- | --- |
| S124 | IGHV2-26 | IGKV1-39 | 15 | Pinto et al. (2020) |
| S309 | IGHV1-18 | IGKV3-20 | 18 | Pinto et al. (2020) |
| S315 | IGHV3-7 | IGLV3-25 | 15 | Pinto et al. (2020) |
| S303 | IGHV3-23 | IGKV1-5 | 15 | Pinto et al. (2020) |
| P1A-1C7 | IGHV1-46 | IGKV1-39 | 13 | Ju et al. (2020) |
| P1A-1C10 | IGHV1-69 | IGKV1-5 | 14 | Ju et al. (2020) |
| P1A-1C11 | IGHV1-69 | IGKV1-5 | 14 | Ju et al. (2020) |
| P1A-1C6 | IGHV3-13 | IGKV1-39 | 17 | Ju et al. (2020) |
| P1A-1D3 | IGHV3-13 | IGKV1-39 | 16 | Ju et al. (2020) |
| P1A-1C2 | IGHV3-23 | IGKV1-36 | 8 | Ju et al. (2020) |
| P1A-1B2 | IGHV3-30 | IGLV2-14 | 10 | Ju et al. (2020) |
| P1A-1C1 | IGHV3-33 | IGKV1D-13 | 15 | Ju et al. (2020) |
| P1A-1D1 | IGHV3-53 | IGLV2-8 | 10 | Ju et al. (2020) |
| P1A-1D5 | IGHV3-53 | IGKV1-33 | 13 | Ju et al. (2020) |
| P1A-1D6 | IGHV3-53 | IGKV1-33 | 13 | Ju et al. (2020) |
| P2A-1A10 | IGHV1-2 | IGKV2-40 | 17 | Ju et al. (2020) |
| P2B-1A4 | IGHV1-2 | IGKV2-40 | 17 | Ju et al. (2020) |
| P2B-1B2 | IGHV1-2 | IGKV2-40 | 17 | Ju et al. (2020) |
| P2B-2G1 | IGHV1-2 | IGKV2-40 | 17 | Ju et al. (2020) |
| P2B-2G12 | IGHV1-2 | IGKV2-40 | 17 | Ju et al. (2020) |
| P2C-1A10 | IGHV1-2 | IGKV2-40 | 17 | Ju et al. (2020) |
| P2C-1B10 | IGHV1-2 | IGKV2-40 | 17 | Ju et al. (2020) |
| P2C-1D6 | IGHV1-2 | IGKV2-40 | 17 | Ju et al. (2020) |
| P2C-1D12 | IGHV1-2 | IGKV2-40 | 17 | Ju et al. (2020) |
| P2C-1F10 | IGHV1-2 | IGKV2-40 | 17 | Ju et al. (2020) |
| P2B-1F8 | IGHV1-2 | IGKV3-20 | 12 | Ju et al. (2020) |
| P2B-2G9 | IGHV1-2 | IGKV3-20 | 12 | Ju et al. (2020) |
| P2B-1C3 | IGHV1-46 | IGKV1-5 | 13 | Ju et al. (2020) |
| P2C-1C10 | IGHV1-69 | IGKV3-11 | 9 | Ju et al. (2020) |
| P2B-2G10 | IGHV1-69 | IGKV1-39 | 9 | Ju et al. (2020) |
| P2B-1F11 | IGHV1-69 | IGLV1-40 | 15 | Ju et al. (2020) |
| P2B-1D9 | IGHV2-5 | IGLV1-47 | 14 | Ju et al. (2020) |
| P2B-1E2 | IGHV2-5 | IGKV1-5 | 10 | Ju et al. (2020) |
| P2B-1E4 | IGHV2-5 | IGLV2-14 | 9 | Ju et al. (2020) |
| P2B-1F4 | IGHV2-70 | IGLV1-44 | 12 | Ju et al. (2020) |
| P2C-1A3 | IGHV3-11 | IGKV1-9 | 10 | Ju et al. (2020) |
| P2B-1D6 | IGHV3-15 | IGLV1-44 | 22 | Ju et al. (2020) |
| P2C-1B12 | IGHV3-15 | IGLV6-57 | 11 | Ju et al. (2020) |
| P2B-1F9 | IGHV3-15 | IGKV1-NL1 | 14 | Ju et al. (2020) |
| P2C-1D5 | IGHV3-23 | IGLV3-21 | 12 | Ju et al. (2020) |
| P2B-1B4 | IGHV3-30 | IGKV1-39 | 20 | Ju et al. (2020) |
| P2B-1F2 | IGHV3-33 | IGLV2-11 | 9 | Ju et al. (2020) |
| P2B-2G4 | IGHV3-33 | IGLV2-11 | 9 | Ju et al. (2020) |
| P2C-1C8 | IGHV3-33 | IGKV2D-30 | 11 | Ju et al. (2020) |
| P2A-1B3 | IGHV3-48 | IGKV3-20 | 14 | Ju et al. (2020) |
| P2B-1B11 | IGHV3-48 | IGKV3-20 | 14 | Ju et al. (2020) |
| P2B-1B12 | IGHV3-48 | IGKV3-20 | 14 | Ju et al. (2020) |
| P2B-1C4 | IGHV3-48 | IGKV3-20 | 14 | Ju et al. (2020) |
| P2B-1E11 | IGHV3-48 | IGKV3-20 | 14 | Ju et al. (2020) |
| P2B-2H7 | IGHV3-48 | IGKV3-20 | 14 | Ju et al. (2020) |
| P2B-1G12 | IGHV3-48 | IGKV3-20 | 14 | Ju et al. (2020) |
| P2C-1E5 | IGHV3-48 | IGKV3-20 | 14 | Ju et al. (2020) |
| P2B-1A10 | IGHV3-53 | IGKV1-33 | 13 | Ju et al. (2020) |
| P2B-1F5 | IGHV3-53 | IGKV1-NL1 | 12 | Ju et al. (2020) |
| P2C-1D7 | IGHV3-53 | IGKV2D-30 | 10 | Ju et al. (2020) |
| P2B-1G1 | IGHV3-66 | IGKV3-20 | 9 | Ju et al. (2020) |
| P2C-1E1 | IGHV3-66 | IGKV3-11 | 7 | Ju et al. (2020) |
| P2C-1F11 | IGHV3-66 | IGKV3-20 | 9 | Ju et al. (2020) |
| P2A-1A8 | IGHV3-9 | IGLV2-14 | 21 | Ju et al. (2020) |
| P2B-1B10 | IGHV3-9 | IGLV2-14 | 21 | Ju et al. (2020) |
| P2B-1C10 | IGHV3-9 | IGLV2-14 | 21 | Ju et al. (2020) |
| P2B-1D3 | IGHV3-9 | IGLV2-14 | 21 | Ju et al. (2020) |
| P2B-2H4 | IGHV3-9 | IGLV2-14 | 21 | Ju et al. (2020) |
| P2C-1A5 | IGHV3-9 | IGLV2-14 | 21 | Ju et al. (2020) |
| P2C-1A8 | IGHV3-9 | IGLV2-14 | 21 | Ju et al. (2020) |
| P2C-1B1 | IGHV3-9 | IGLV2-14 | 21 | Ju et al. (2020) |
| P2C-1C12 | IGHV3-9 | IGLV2-14 | 21 | Ju et al. (2020) |
| P2C-1A6 | IGHV3-9 | IGLV2-14 | 21 | Ju et al. (2020) |
| P2A-1A9 | IGHV3-9 | IGLV1-40 | 15 | Ju et al. (2020) |
| P2C-1A1 | IGHV3-9 | IGLV1-40 | 15 | Ju et al. (2020) |
| P2B-2G11 | IGHV3-9 | IGLV1-40 | 15 | Ju et al. (2020) |
| P2B-1E12 | IGHV3-9 | IGLV3-20 | 15 | Ju et al. (2020) |
| P2B-2F6 | IGHV4-38 | IGLV2-8 | 18 | Ju et al. (2020) |
| P2A-1B10 | IGHV4-39 | IGLV1-47 | 18 | Ju et al. (2020) |
| P2B-1B9 | IGHV4-39 | IGKV1-NL1 | 7 | Ju et al. (2020) |
| P2B-2F11 | IGHV4-39 | IGKV1-NL1 | 7 | Ju et al. (2020) |
| P2B-1G8 | IGHV4-39 | IGKV1-5 | 9 | Ju et al. (2020) |
| P2B-1A1 | IGHV4-59 | IGLV2-14 | 12 | Ju et al. (2020) |
| P2B-1D11 | IGHV4-59 | IGLV3-25 | 20 | Ju et al. (2020) |
| P2B-1F11 | IGHV4-59 | IGKV1-39 | 13 | Ju et al. (2020) |
| P2C-1A7 | IGHV5-51 | IGLV3-1 | 15 | Ju et al. (2020) |
| P2B-1A12 | IGHV7-4-1 | IGKV1-39 | 14 | Ju et al. (2020) |
| P2B-1G5 | IGHV7-4-1 | IGLV3-21 | 10 | Ju et al. (2020) |
| P3A-1F1 | IGHV3-13 | IGKV1-39 | 15 | Ju et al. (2020) |
| P3A-1G8 | IGHV3-64 | IGLV1-44 | 17 | Ju et al. (2020) |
| P4A-2A10 | IGHV1-46 | IGLV1-40 | 24 | Ju et al. (2020) |
| P4B-1F6 | IGHV1-69 | IGLV2-23 | 13 | Ju et al. (2020) |
| P4B-1E11 | IGHV2-5 | IGLV1-36 | 16 | Ju et al. (2020) |
| P4A-2A2 | IGHV3-23 | IGLV1-51 | 12 | Ju et al. (2020) |
| P4A-2A8 | IGHV3-23 | IGLV3-21 | 9 | Ju et al. (2020) |
| P4A-2C1 | IGHV3-23 | IGKV2-28 | 14 | Ju et al. (2020) |
| P4A-1H5 | IGHV3-30 | IGKV1-39 | 19 | Ju et al. (2020) |
| P4B-1G2 | IGHV3-30 | IGKV1-39 | 19 | Ju et al. (2020) |
| P4A-2B3 | IGHV3-30 | IGKV1-39 | 19 | Ju et al. (2020) |
| P4A-1H6 | IGHV3-30 | IGKV1-39 | 19 | Ju et al. (2020) |
| P4B-1G5 | IGHV3-30 | IGLV3-21 | 20 | Ju et al. (2020) |
| P4A-2E10 | IGHV3-30 | IGKV1-39 | 19 | Ju et al. (2020) |
| P4B-1E3 | IGHV3-30 | IGKV1-39 | 19 | Ju et al. (2020) |
| P4A-2D9 | IGHV3-30 | IGKV1-39 | 19 | Ju et al. (2020) |
| P4B-1F4 | IGHV3-30 | IGKV2-30 | 20 | Ju et al. (2020) |
| P4B-1E7 | IGHV3-43D | IGKV3-1 | 18 | Ju et al. (2020) |
| P4B-1F10 | IGHV3-7 | IGKV3-21 | 11 | Ju et al. (2020) |
| P4A-2D1 | IGHV3-9 | IGKV1-12 | 11 | Ju et al. (2020) |
| P4A-2D2 | IGHV4-39 | IGKV3-20 | 14 | Ju et al. (2020) |
| P4B-1E12 | IGHV4-59 | IGLV1-44 | 9 | Ju et al. (2020) |
| P4A-2C12 | IGHV5-51 | IGLV1-44 | 13 | Ju et al. (2020) |
| P8A-1A8 | IGHV3-23 | IGLV3-21 | 9 | Ju et al. (2020) |
| P8A-1C6 | IGHV3-30 | IGKV1-33 | 18 | Ju et al. (2020) |
| P8A-1A5 | IGHV5-51 | IGLV1-47 | 16 | Ju et al. (2020) |
| P8A-1D5 | IGHV6-1 | IGKV3-20 | 14 | Ju et al. (2020) |
| P5A-1A1 | IGHV1-24 | IGKV2-28 | 13 | Ju et al. (2020) |
| P5A-1C8 | IGHV1-46 | IGKV1-33 | 20 | Ju et al. (2020) |
| P5A-2D5 | IGHV1-46 | IGLV1-40 | 22 | Ju et al. (2020) |
| P5A-2C8 | IGHV1-46 | IGLV2-23 | 13 | Ju et al. (2020) |
| P5A-2E9 | IGHV1-46 | IGLV2-14 | 20 | Ju et al. (2020) |
| P5A-3B8 | IGHV1-46 | IGLV2-23 | 14 | Ju et al. (2020) |
| P5A-3A11 | IGHV1-69 | IGKV1-39 | 12 | Ju et al. (2020) |
| P5A-3C10 | IGHV1-69 | IGLV6-57 | 20 | Ju et al. (2020) |
| P5A-1A2 | IGHV1-8 | IGLV1-40 | 19 | Ju et al. (2020) |
| P5A-1C11 | IGHV1-8 | IGLV3-21 | 15 | Ju et al. (2020) |
| P5A-2F11 | IGHV1-8 | IGKV4-1 | 13 | Ju et al. (2020) |
| P5A-3B9 | IGHV1-8 | IGKV1-36 | 13 | Ju et al. (2020) |
| P5A-2C12 | IGHV2-5 | IGKV3-11 | 14 | Ju et al. (2020) |
| P5A-3C12 | IGHV2-5 | IGKV4-1 | 17 | Ju et al. (2020) |
| P5A-3C3 | IGHV2-5 | IGLV6-57 | 10 | Ju et al. (2020) |
| P5A-3C1 | IGHV3-11 | IGLV3-21 | 11 | Ju et al. (2020) |
| P5A-1C4 | IGHV3-13 | IGKV1-39 | 18 | Ju et al. (2020) |
| P5A-2G8 | IGHV3-13 | IGKV1-39 | 11 | Ju et al. (2020) |
| P5A-2D3 | IGHV3-13 | IGKV1-39 | 14 | Ju et al. (2020) |
| P5A-3B10 | IGHV3-13 | IGKV1-39 | 14 | Ju et al. (2020) |
| P5A-1D8 | IGHV3-13 | IGLV3-19 | 16 | Ju et al. (2020) |
| P5A-2G10 | IGHV3-13 | IGLV3-19 | 16 | Ju et al. (2020) |
| P5A-2H6 | IGHV3-15 | IGLV3-19 | 16 | Ju et al. (2020) |
| P5A-1D6 | IGHV3-23 | IGLV3-21 | 11 | Ju et al. (2020) |
| P5A-2E12 | IGHV3-23 | IGLV3-21 | 12 | Ju et al. (2020) |
| P5A-3D12 | IGHV3-23 | IGLV1-47 | 22 | Ju et al. (2020) |
| P5A-1B6 | IGHV3-30 | IGKV1-33 | 18 | Ju et al. (2020) |
| P5A-2E6 | IGHV3-30 | IGKV1-33 | 18 | Ju et al. (2020) |
| P5A-1B1 | IGHV3-33 | IGKV3-15 | 12 | Ju et al. (2020) |
| P5A-1C5 | IGHV3-33 | IGKV3-15 | 12 | Ju et al. (2020) |
| P5A-2H7 | IGHV3-33 | IGKV3-15 | 12 | Ju et al. (2020) |
| P5A-2G9 | IGHV3-33 | IGLV5-37 | 10 | Ju et al. (2020) |
| P5A-2G11 | IGHV3-33 | IGLV2-14 | 15 | Ju et al. (2020) |
| P5A-1B8 | IGHV3-53 | IGKV1-9 | 7 | Ju et al. (2020) |
| P5A-1D2 | IGHV3-53 | IGLV1-40 | 13 | Ju et al. (2020) |
| P5A-1D1 | IGHV3-53 | IGKV1-9 | 9 | Ju et al. (2020) |
| P5A-2C9 | IGHV3-7 | IGKV3-20 | 12 | Ju et al. (2020) |
| P5A-2E4 | IGHV3-7 | IGKV3-20 | 12 | Ju et al. (2020) |
| P5A-2G12 | IGHV3-7 | IGLV6-57 | 10 | Ju et al. (2020) |
| P5A-2D12 | IGHV3-7 | IGKV2-28 | 16 | Ju et al. (2020) |
| P5A-2F1 | IGHV3-74 | IGLV6-57 | 10 | Ju et al. (2020) |
| P5A-1C10 | IGHV3-9 | IGLV3-21 | 12 | Ju et al. (2020) |
| P5A-2E8 | IGHV3-9 | IGLV3-21 | 11 | Ju et al. (2020) |
| P5A-3A2 | IGHV3-9 | IGLV3-21 | 12 | Ju et al. (2020) |
| P5A-2D6 | IGHV3-9 | IGLV1-40 | 12 | Ju et al. (2020) |
| P5A-1B12 | IGHV3-9 | IGLV1-51 | 15 | Ju et al. (2020) |
| P5A-3A6 | IGHV3-9 | IGLV2-14 | 25 | Ju et al. (2020) |
| P5A-3D9 | IGHV3-9 | IGKV3-15 | 14 | Ju et al. (2020) |
| P5A-1D10 | IGHV3-11 | IGLV2-14 | 19 | Ju et al. (2020) |
| P5A-3A1 | IGHV3-53 | IGKV3-20 | 9 | Ju et al. (2020) |
| P5A-3C8 | IGHV3-53 | IGKV1-9 | 9 | Ju et al. (2020) |
| P5A-2D10 | IGHV4-31 | IGLV6-57 | 10 | Ju et al. (2020) |
| P5A-2G5 | IGHV4-31 | IGLV3-21 | 12 | Ju et al. (2020) |
| P5A-1A12 | IGHV4-39 | IGKV4-1 | 15 | Ju et al. (2020) |
| P5A-2C7 | IGHV4-39 | IGLV2-23 | 14 | Ju et al. (2020) |
| P5A-2F7 | IGHV4-39 | IGLV2-23 | 16 | Ju et al. (2020) |
| P5A-2F9 | IGHV4-39 | IGLV2-23 | 12 | Ju et al. (2020) |
| P5A-1A5 | IGHV4-4 | IGLV2-14 | 12 | Ju et al. (2020) |
| P5A-1C6 | IGHV4-4 | IGLV1-40 | 20 | Ju et al. (2020) |
| P5A-3A10 | IGHV4-4 | IGKV1-39 | 19 | Ju et al. (2020) |
| P5A-1B9 | IGHV4-59 | IGKV4-1 | 20 | Ju et al. (2020) |
| P5A-3A7 | IGHV4-59 | IGKV4-1 | 20 | Ju et al. (2020) |
| P5A-3B1 | IGHV4-59 | IGKV4-1 | 20 | Ju et al. (2020) |
| P5A-3B6 | IGHV4-59 | IGKV4-1 | 20 | Ju et al. (2020) |
| P5A-2C10 | IGHV4-59 | IGLV3-21 | 15 | Ju et al. (2020) |
| P5A-2E5 | IGHV4-59 | IGLV6-57 | 10 | Ju et al. (2020) |
| P5A-2G4 | IGHV4-59 | IGKV1D-16 | 10 | Ju et al. (2020) |
| P5A-2G7 | IGHV4-61 | IGLV2-14 | 18 | Ju et al. (2020) |
| P5A-1B10 | IGHV5-51 | IGKV2-28 | 10 | Ju et al. (2020) |
| P5A-1C9 | IGHV5-51 | IGLV3-19 | 9 | Ju et al. (2020) |
| P5A-2D11 | IGHV5-51 | IGLV1-44 | 11 | Ju et al. (2020) |
| P5A-3B4 | IGHV5-51 | IGLV1-44 | 11 | Ju et al. (2020) |
| P5A-2H3 | IGHV5-51 | IGLV1-44 | 11 | Ju et al. (2020) |
| P5A-2E1 | IGHV5-51 | IGLV3-21 | 10 | Ju et al. (2020) |
| P5A-1B11 | IGHV7-4-1 | IGKV1-39 | 18 | Ju et al. (2020) |
| P5A-2D7 | IGHV7-4-1 | IGKV6-21 | 8 | Ju et al. (2020) |
| P5A-3C9 | IGHV7-4-1 | IGKV6-21 | 8 | Ju et al. (2020) |
| P5A-3D11 | IGHV7-4-1 | IGKV6-21 | 8 | Ju et al. (2020) |
| P16A-1A3 | IGHV1-3 | IGLV6-57 | 9 | Ju et al. (2020) |
| P16A-1A8 | IGHV1-46 | IGLV3-21 | 18 | Ju et al. (2020) |
| P16A-1B5 | IGHV1-46 | IGLV3-21 | 11 | Ju et al. (2020) |
| P16A-1C6 | IGHV1-46 | IGLV3-21 | 14 | Ju et al. (2020) |
| P16A-1C1 | IGHV3-13 | IGKV1-39 | 19 | Ju et al. (2020) |
| P16A-1A5 | IGHV3-33 | IGKV1-33 | 13 | Ju et al. (2020) |
| P16A-1A12 | IGHV3-33 | IGLV1-51 | 17 | Ju et al. (2020) |
| P16A-1B1 | IGHV3-74 | IGLV1-36 | 13 | Ju et al. (2020) |
| P16A-1B3 | IGHV3-9 | IGLV3-1 | 22 | Ju et al. (2020) |
| P16A-1B12 | IGHV4-34 | IGLV1-51 | 14 | Ju et al. (2020) |
| P16A-1B8 | IGHV5-51 | IGLV3-1 | 17 | Ju et al. (2020) |
| P16A-1A7 | IGHV7-4-1 | IGLV3-21 | 12 | Ju et al. (2020) |
| P16A-1A10 | IGHV7-4-1 | IGLV3-21 | 13 | Ju et al. (2020) |
| P22A-1E10 | IGHV1-46 | IGKV3-11 | 13 | Ju et al. (2020) |
| P22A-1D2 | IGHV1-8 | IGLV1-40 | 19 | Ju et al. (2020) |
| P22A-1D8 | IGHV3-23 | IGKV3-15 | 18 | Ju et al. (2020) |
| P22A-1D7 | IGHV3-33 | IGKV1-39 | 11 | Ju et al. (2020) |
| P22A-1D1 | IGHV3-53 | IGKV1-9 | 9 | Ju et al. (2020) |
| P22A-1E8 | IGHV3-9 | IGKV3-15 | 14 | Ju et al. (2020) |
| P22A-1D5 | IGHV4-39 | IGLV2-23 | 12 | Ju et al. (2020) |
| P22A-1E6 | IGHV4-59 | IGKV3-20 | 14 | Ju et al. (2020) |
| BD-494 | IGHV3-53 | IGKV1-9 | 9 | Cao et al. (2020) |
| BD-498 | IGHV3-66 | IGKV1-9 | 9 | Cao et al. (2020) |
| BD-500 | IGHV3-53 | IGKV1D-39 | 9 | Cao et al. (2020) |
| BD-503 | IGHV3-53 | IGKV1D-39 | 9 | Cao et al. (2020) |
| BD-504 | IGHV3-66 | IGKV1-9 | 9 | Cao et al. (2020) |
| BD-505 | IGHV3-53 | IGKV1D-33 | 9 | Cao et al. (2020) |
| BD-506 | IGHV3-53 | IGKV1-9 | 9 | Cao et al. (2020) |
| BD-507 | IGHV3-53 | IGKV1-9 | 9 | Cao et al. (2020) |
| BD-508 | IGHV3-53 | IGKV1D-39 | 9 | Cao et al. (2020) |
| CR3022 | IGHV5-51 | IGKV4-1 | 10 | Yuan et al. (2020) |
| CC12.1 | IGHV3-53 | IGKV1-9 | - | Rogers et al. (2020) |
| CC12.2 | IGHV3-53 | IGKV3-20 | - | Rogers et al. (2020) |
| CC12.3 | IGHV3-53 | IGKV3-20 | - | Rogers et al. (2020) |
| CC12.4 | IGHV1-2 | IGLV2-8 | - | Rogers et al. (2020) |
| CC12.5 | IGHV1-2 | IGLV2-14 | - | Rogers et al. (2020) |
| CC12.6 | IGHV1-2 | IGLV2-14 | - | Rogers et al. (2020) |
| CC12.7 | IGHV1-2 | IGLV2-14 | - | Rogers et al. (2020) |
| CC12.8 | IGHV1-2 | IGLV2-14 | - | Rogers et al. (2020) |
| CC12.9 | IGHV1-2 | IGLV2-14 | - | Rogers et al. (2020) |
| CC12.10 | IGHV1-2 | IGLV2-14 | - | Rogers et al. (2020) |
| CC12.11 | IGHV1-2 | IGLV2-14 | - | Rogers et al. (2020) |
| CC12.12 | IGHV1-2 | IGLV2-14 | - | Rogers et al. (2020) |
| CC12.13 | IGHV3-53 | IGKV1-33 | - | Rogers et al. (2020) |
| CC12.14 | IGHV3-21 | IGKV2-30 | - | Rogers et al. (2020) |
| CC12.15 | IGHV3-48 | IGLV1-40 | - | Rogers et al. (2020) |
| CC12.16 | IGHV3-33 | IGLV3-21 | - | Rogers et al. (2020) |
| CC12.17 | IGHV3-30 | IGLV3-21 | - | Rogers et al. (2020) |
| CC12.18 | IGHV1-46 | IGLV6-57 | - | Rogers et al. (2020) |
| CC12.19 | IGHV3-23 | IGLV3-21 | - | Rogers et al. (2020) |
| COVA1-07 | IGHV1-69 | - | 13 | Brouwer et al. (2020) |
| COVA1-08 | IGHV3-30 | - | 12 | Brouwer et al. (2020) |
| COVA1-10 | IGHV3-66 | - | 19 | Brouwer et al. (2020) |
| COVA1-12 | IGHV1-2 | - | 13 | Brouwer et al. (2020) |
| COVA1-16 | IGHV1-46 | - | 20 | Brouwer et al. (2020) |
| COVA1-18 | IGHV3-66 | - | 10 | Brouwer et al. (2020) |
| COVA2-01 | IGHV3-13 | - | 12 | Brouwer et al. (2020) |
| COVA2-02 | IGHV4-39 | - | 13 | Brouwer et al. (2020) |
| COVA2-04 | IGHV3-53 | - | 10 | Brouwer et al. (2020) |
| COVA2-05 | IGHV5-51 | - | 18 | Brouwer et al. (2020) |
| COVA2-07 | IGHV3-53 | - | 7 | Brouwer et al. (2020) |
| COVA2-11 | IGHV3-21 | - | 17 | Brouwer et al. (2020) |
| COVA2-13 | IGHV1-69 | - | 10 | Brouwer et al. (2020) |
| COVA2-15 | IGHV3-23 | - | 20 | Brouwer et al. (2020) |
| COVA2-16 | IGHV1-69 | - | 14 | Brouwer et al. (2020) |
| COVA2-17 | IGHV1-69 | - | 11 | Brouwer et al. (2020) |
| COVA2-20 | IGHV3-53 | - | 15 | Brouwer et al. (2020) |
| COVA2-23 | IGHV1-2 | - | 18 | Brouwer et al. (2020) |
| COVA2-24 | IGHV5-10 | - | 18 | Brouwer et al. (2020) |
| COVA2-27 | IGHV1-8 | - | 14 | Brouwer et al. (2020) |
| COVA2-29 | IGHV4-30 | - | 18 | Brouwer et al. (2020) |
| COVA2-31 | IGHV1-2 | - | 16 | Brouwer et al. (2020) |
| COVA2-32 | IGHV1-69 | - | 13 | Brouwer et al. (2020) |
| COVA2-36 | IGHV5-51 | - | 14 | Brouwer et al. (2020) |
| COVA2-39 | IGHV3-53 | - | 15 | Brouwer et al. (2020) |
| COVA2-44 | IGHV3-30 | - | 13 | Brouwer et al. (2020) |
| COVA2-45 | IGHV1-2 | - | 22 | Brouwer et al. (2020) |
| COVA2-46 | IGHV4-39 | - | 10 | Brouwer et al. (2020) |
| COVA3-05 | IGHV1-24 | - | 14 | Brouwer et al. (2020) |
| COVA3-06 | IGHV1-69 | - | 16 | Brouwer et al. (2020) |
| COVA3-09 | IGHV4-59 | - | 12 | Brouwer et al. (2020) |
| COVA3-10 | IGHV5-51 | - | 14 | Brouwer et al. (2020) |
| B5 | IGHV1-2 | IGKV3-20 | - | Wu et al. (2020) |
| B38 | IGHV3-53 | IGKV1-9 | 7 | Wu et al. (2020) |
| H2 | IGHV3-9 | IGKV1-39 | - | Wu et al. (2020) |
| H4 | IGHV1-2 | IGKV2-40 | 17 | Wu et al. (2020) |
| COV21.1 | IGHV1-58 | IGKV3-20 | 14 | Robbiani et al. (2020) |
| COV21.2 | IGHV1-58 | IGKV3-20 | 14 | Robbiani et al. (2020) |
| COV57.1 | IGHV1-58 | IGKV3-20 | 14 | Robbiani et al. (2020) |
| COV57.2 | IGHV1-58 | IGKV3-20 | 14 | Robbiani et al. (2020) |
| COV107.1 | IGHV1-58 | IGKV3-20 | 14 | Robbiani et al. (2020) |
| COV107.2 | IGHV1-58 | IGKV3-20 | 14 | Robbiani et al. (2020) |
| COV21.3 | IGHV3-30 | IGKV1-39 | 14 | Robbiani et al. (2020) |
| COV21.4 | IGHV3-30 | IGKV1-39 | 12 | Robbiani et al. (2020) |
| COV72.1 | IGHV3-30 | IGKV1-39 | 14 | Robbiani et al. (2020) |
| COV72.2 | IGHV3-30 | IGKV1-39 | 14 | Robbiani et al. (2020) |
| COV72.3 | IGHV3-30 | IGKV1-39 | 14 | Robbiani et al. (2020) |
| 1M-1D2 | IGHV3-64 | IGLV1-47 | 18 | Chi et al. (2020) |
| 2M-10B11 | IGHV3-66 | IGLV6-57 | 10 | Chi et al. (2020) |
| 2M-4G4 | IGHV1-46 | IGLV2-23 | 23 | Chi et al. (2020) |
| CV5 | IGHV1-46 | IGKV4-1 | - | Seydoux et al. (2020) |
| CV30 | IGHV3-53 | IGKV3-20 | - | Seydoux et al. (2020) |
| CV43 | IGHV3-30 | IGLV6-57 | - | Seydoux et al. (2020) |
| CA1 | IGHV1-18 | IGKV3-11 | 21 | Shi et al. (2020) |
| CB6 | IGHV3-66 | IGKV1-39 | 11 | Shi et al. (2020) |
| C105 | IGHV3-53 | IGLV2-8 | - | Barnes et al. (2020) |

**Table S3.** Hydrogen bonds and salt bridges identified at the antibody-RBD interface using the PISA program.

| SARS-CoV-2 RBD | Distance [Å] | CC12.1 |
| --- | --- | --- |
| Hydrogen bonds |  |  |
| TYR 473[OH] | 3.63 | VH SER 53[OG] |
| TYR 473[OH] | 2.77 | VH SER 31[O] |
| ASP 420[OD2] | 2.51 | VH SER 56[OG] |
| TYR 421[O] | 3.39 | VH SER 53[N] |
| LEU 455[O] | 2.59 | VH TYR 33[OH] |
| ALA 475[O] | 3.02 | VH ASN 32[ND2] |
| ALA 475[O] | 3.16 | VH THR 28[N] |
| ASN 487[OD1] | 3.13 | VH ARG 94[NH1] |
| ASN 487[OD1] | 3.16 | VH ARG 94[NH2] |
| TYR 489[OH] | 3.26 | VH ARG 94[NH2] |
| GLN 493[OE1] | 2.85 | VH TYR 99[OH] |
| TYR 505[OH] | 2.95 | VL LEU 91[O] |
| ARG 403[NH2] | 2.08 | VL ASN 92[O] |
| TYR 505[OH] | 3.11 | VL ASN 92[O] |
| SER 494[O] | 3.55 | VL TYR 32[OH] |
| GLN 498[OE1] | 3.61 | VL SER 67[OG] |
| TYR 505[OH] | 2.94 | VL GLN 90[NE2] |
| TYR 453[OH] | 3.29 | VL ASN 92[ND2] |
| THR 415[O] | 2.69 | VL TYR 94[OH] |
| Salt bridges |  |  |
| LYS 417[NZ] | 3.17 | VH ASP 97[OD1] |

| SARS-CoV-2 RBD | Distance [Å] | CC12.3 |
| --- | --- | --- |
| Hydrogen bonds |  |  |
| ASP 420[OD2] | 2.60 | VH SER 56[OG] |
| TYR 421[OH] | 3.36 | VH SER 53[OG] |
| TYR 421[OH] | 3.72 | VH SER 53[N] |
| LEU 455[O] | 2.65 | VH TYR 33[OH] |
| ARG 457[O] | 2.81 | VH SER 53[OG] |
| ALA 475[O] | 2.97 | VH ASN 32[ND2] |
| ALA 475[O] | 3.13 | VH THR 28[N] |
| ASN 487[OD1] | 2.67 | VH ARG 94[NH2] |
| TYR 489[OH] | 2.88 | VH ARG 94[NH1] |
| TYR 489[OH] | 2.71 | VH ARG 94[NH2] |
| SER 477[N] | 3.88 | VH THR 28[OG1] |
| TYR 473[OH] | 2.74 | VH SER 31[O] |
| ARG 457[N] | 3.64 | VL SER 53[OG] |
| TYR 495[O] | 3.87 | VL TYR 32[OH] |
| TYR 505[OH] | 3.88 | VL SER 93[N] |
| TYR 505[OH] | 3.21 | VL SER 28[O] |

